# A novel multi-epitope peptide vaccine candidate targeting Hepatitis E virus: an in-silico approach

**DOI:** 10.1101/2022.12.13.520355

**Authors:** Anoop Kumar, Utkarsha Sahu, Geetanjali Agnihotri, Anshuman Dixit, Prashant Khare

## Abstract

HEV is a foodborne virus transmitted through the fecal-oral route that causes viral hepatitis in humans worldwide. Ever since its discovery as a zoonotic agent, HEV was isolated from several species with an expanding range of hosts. HEV possesses several features of other RNA viruses but also has certain HEV-specific traits that make its viral-host interactions inimitable. HEV leads to severe morbidity and mortality in immunocompromised people and pregnant women across the world. The situation in underdeveloped countries is even more alarming. Even after creating a menace across the world, we still lack an effective vaccine against HEV. Till date, there is only one licensed vaccine for HEV available only in China. The development of an anti-HEV vaccine that can reduce HEV-induced morbidity and mortality is required. Live attenuated and killed vaccines against HEV are not accessible due to the lack of a tolerant cell culture system, slow viral replication kinetics and varying growth conditions. Thus, the main focus for anti-HEV vaccine development is now on the molecular approaches. In the current study, we have designed a multi-epitope vaccine against HEV through a reverse vaccinology approach **(Figure 1)**. For the first time, we have used viral ORF3, capsid protein and polyprotein altogether for epitope prediction. These are crucial for viral replication and persistence and are major vaccine targets against HEV. The proposed *in-silico* vaccine construct comprises of highly immunogenic and antigenic T-cell and B-cell epitopes of HEV proteins and ORF3. The construct is capable of inducing an effective and long-lasting host immune response as evident from the simulations results. In addition, the construct is stable, non-allergic and antigenic for the host. Altogether, our findings suggest that the *in-silico* vaccine construct may be useful as a vaccine candidate for preventing HEV infections.

## 1. Introduction

Hepatitis E virus (HEV) is a positive sense single-stranded RNA (**+**ssRNA) virus belonging to the *Hepeviridae* family (1). HEV is the leading cause of virus-induced acute hepatitis in humans worldwide. Approximately, 35 million cases of HEV infection and about 70, 000 deaths are reported annually across the world (2). The average mortality rate due to HEV infection is around 0.2-4%, while in pregnant women this risk increases up to 10-25% (3). The underdeveloped countries from the Indian subcontinent and Africa are most affected by HEV-induced hepatitis (4). In underdeveloped and developing countries the lack of proper sanitation facilities leads to approximately 3.3 million HEV cases and about 56,600 deaths every year (3,5).

Even though most HEV infections are self-limiting, there is still a subpopulation of individuals such as pregnant women and immunocompromised people at a high risk of HEV infection with a mortality rate of up to 30% (6). A vaccine (Hecolin^®^) is presently approved for administration only in China, not in the rest of the countries across the world (7). Currently, available therapeutics against HEV such as Interferon-α and ribavirin were reported to possess serious side effects, and treatment failure and are also contraindicated for pregnant women (8). Also, around 20 direct-acting antiviral agents (DAAs) are under phase II and phase III clinical trials but their efficacy against HEV is not well established since most of them were initially designed for HCV proteins (9). Altogether, there is an urgent need for effective and safer treatment options such as vaccines and antivirals to limit severe HEV infection in the target population.

HEV is a small virus of about 320–340 Å particle size and a 7200bp long genome (10). The HEV genome codes for three open reading frames (ORFs)-ORF1, ORF2 and ORF3. In addition, a 5′ 7-methylguanosine cap, short 5′ untranslated region (UTR) and a 3′UTR region are also present (11). The ORF1 of the HEV genome codes for a polyprotein with different functional domains such as helicase (Hel), papain-like cysteine protease (PCP), RNA-dependent RNA polymerase (RdRp), methyltransferase (Met), proline-rich region (PRR) or hypervariable region (HVR), X and Y domain (12,13). ORF2 encodes for the viral capsid (14) and ORF3 for a multifunctional phosphoprotein known as VP13 (11). Thus, these proteins and/or ORFs are crucial targets for vaccine construct against HEV. HEV has four different genotypes having varying host range and geographical distributions (15). Regardless of the different genotypes, the HEV strains possess a single serotype and cross-reactive epitopes because of high sequence conservancy (15).

In the current study, we have undertaken the designing of a multiepitope vaccine against HEV via *in-silico* approach. The current vaccine construct comprises highly immunogenic and antigenic MHC I and MHC II epitopes of the major viral proteins and ORF. The epitopes were linked using suitable linkers and an adjuvant for added for further enhancement of the effectiveness of the construct. Once the construct sequence was prepared the secondary and tertiary structures were predicted. The construct was docked with toll-like receptor (TLR)3 and TLR4 which indicated stable interaction with host TLRs. Moreover, the vaccine construct was cloned and further validated using *in-silico* immune simulation tools. The results indicated that the construct had stable interactions and was capable of inducing a prolonged and effective immune response in the host. However, further *in vitro* and *in vivo* experiments are required to establish the validity of the construct. Altogether, this construct employs the reverse vaccinology approach and represents a novel *in-silico* vaccine construct. It would further pave the way for the development of effective HEV vaccines in the future helping the target population comprising mainly of pregnant women, immunocompromised individuals and populations from under-developed countries.

## 2. Material and Methods

### 2.1. Protein sequences

The sequence of HEV capsid protein, polyprotein and ORF3, were extracted from the National Center for Biotechnology Information (https://www.ncbi.nlm.nih.gov/).

### 2.2. Prediction of T-cell epitopes

The Immune Epitope Database (IEDB) server was used to predict the MHC-I and MHC-II epitopes for protein sequences of capsid protein, ORF3 & polyprotein of HEV using the default parameters. The percentile score (lower the percentile value higher the binding affinity of the predicted epitope) was chosen <0.5 for MHC-I alleles and <1 for MHC-II alleles

### 2.3. Prediction of Immunogenicity of MHC Class I epitopes

The MHC I epitopes identified from IEDB server were further investigated for their immunogenicity by using IEDB MHC Class I Immunogenicity tool (http://tools.iedb.org/immunogenicity/) that only scrutinized and validated on 9 mer peptides.

### 2.4. Prediction of Antigenicity & Interferon-γ inducing epitope

The antigenicity of the predicted epitopes was evaluated using Vaxijen online tool (16). To analyze the interferon-gamma (IFN-γ) inducing ability of predicted epitopes IFNepitope server was used (https://crdd.osdd.net/raghava/ifnepitope/predict.php) (16). In this analysis, we did Motif and SVM hybrid algorithms (accuracy of %) and IFN-γ versus non-IFN-γ model prediction (17).

### 2.5. Population coverage analysis

The MHC HLA allele variation occurs among various geographical regions across the world. Thus, population coverage analysis becomes obligatory for designing an operative vaccine construct. To evaluate the worldwide coverage of MHC -I and MHC-II allele interacting epitopes, population coverage analysis was done using the IEDB population coverage analysis tool (http://tools.iedb.org/population/). Most potential predominant epitopes were identified from different parts of the world.

### 2.6. Preparation of vaccine construct and analysis of its physiochemical properties

An *in-silico* vaccine construct comprising of MHC I and MHC II epitopes from HEV was constructed by joining the MHC I epitopes with AAY linker and MHC II epitopes with the GPGPG linkers. In addition, the predicted construct has a β-defensin adjuvant at the N terminal. The adjuvant β-defensin is previously reported to be used in diverse *in-silico* vaccine constructs (18) and viral infections (19). Finally, the physiochemical properties of the *in-silico* construct were investigated via the Protparam server.

### 2.7. B-cell epitope Prediction

B-cell epitopes are the main antigenic determinants identified by the host immune system and characterize specific antigenic portion that binds with the B lymphocytes. Initiation of an effective B-cell humoral immune response is necessary for a vaccine candidate. We carried out the artificial neural network-based ABCpred server (http://www.imtech.res.in/raghava/abcpred/ABCsubmission.html) analysis for the prediction of B cell epitopes using default parameters and more than 0.8 epitope score (20).

### 2.8. Prediction of allergenicity of the epitopes

For the prediction of allergenicity of peptides, AllerTOP v2.0 server was used (https://www.ddg-pharmfac.net/AllerTOP/) that employs the auto cross-covariance (ACC) technique.

### 2.9. Prediction of Secondary and tertiary structure

Psipred online server was used to predict the secondary structure of the prepared multi-epitope vaccine construct (http://bioinf.cs.ucl.ac.uk/psipred/). I-TASSER server was used to predict the tertiary structure of the construct (https://zhanglab.ccmb.med.umich.edu/I-TASSER/) (21).

### 2.10. Validation and Refinement of the Tertiary structure

The validation of 3D structure of the prepared construct is important to check for potential errors in the 3D structure. The tertiary structure of the vaccine construct was validated by PDBsum pictorial database that generated a Ramachandran plot giving an imprint of 3D structure contents in PDB (http://www.ebi.ac.uk/thornton-srv/databases/pdbsum/Generate.html).

### 2.11. Molecular docking of multi epitope vaccine construct with TLR3 and TLR4

The multi epitope vaccine construct was docked with TLR3 (2A0Z) and TLR4 (3FXI) to predict its efficiency using the PatchDock online server (https://bioinfo3d.cs.tau.ac.il/PatchDock/php.php). PatchDock is a fast and effective server that operates on specific algorithms. The refinement and re-scoring of docking results were carried out by FireDock server (http://bioinfo3d.cs.tau.ac.il/FireDock/php.php) which gives better results with the least global energy. At last, the model with the best docking score was used and visualized by UCSF Chimera tool for further analysis (22).

### 2.12. MD Simulation analysis

VMD version 1.93 was utilized to generate the preliminary topology and coordinates for all protein complexes. The complexes were solvated in a rectangular water box using a TIP3P water model. A buffering distance of 10 Å was kept. To ensure the electro-neutrality of the solvated system counterions were added to the system. The water molecules were handled using the SETLE algorithm. In the end, the charmm34 force field was used to create topology and coordinates for the simulation. NAMD version 1.9 was used to run the simulations. A stepwise minimization protocol was used to minimize each system prior to simulation. The first step was to minimize the water molecules and the ions while keeping the protein fixed. It was followed by side chains and hydrogen atoms of the complex (100000 steps). Afterward, energy minimization was applied to the entire system for 100000 steps, with small restraints on Cα atoms and DNA backbone atoms (10 kcal/mol). As part of the equilibration protocol, the system was heated from 0 to 310 K in 30 K increments using a canonical ensemble (NVT). Each step of the simulation was run for 20 picoseconds (ps) to allow the system to adapt to the temperature. An isobaric and isothermic ensemble (NPT) with a constant pressure of 1.0 bar was applied at 310 K for 100 ps. After removing the restraints, the system was equilibrated for 1 ns using Langevin piston coupling. Long-range electrostatic interactions were handled using the Particle Mesh Ewald sum algorithm (PME). The SHAKE algorithm was used to constrain hydrogens. As a final step, a 100 ns production run was carried out. Analysis of MD trajectories was conducted to understand complex structure and dynamics. By using VMD, the simulation trajectories were analyzed for root mean square deviation (RMSD), root mean square fluctuation (RMSF), hydrogen bonds, and salt bridges.

### 2.13. *In-silico* cloning optimization of designed vaccine candidate

JCAT (http://www.prodoric.de/JCat) was used to optimize codons and analyze reverse translations of the multi-epitope vaccine candidate. A codon adaptation index (CAI) was used to determine the level of protein expression. Moreover, JCat output provided the GC content percentage (Grote et al., 2005). In the following steps, the optimized sequence has been cloned into pcDNA3.1/V5/His-TOPO/LacZ vector using DNASTAR (https://www.dnastar.com). The default parameters of the server were used for 50 simulation volumes, 1000 steps and 3 injections of our vaccine construct within the 4 weeks interval.

## 3. Results

### 3.1 Prediction of T Lymphocyte Epitopes

The cytotoxic T lymphocytes (CTL) and helper T lymphocytes (HTL) play a vital role in generating a pro-long adaptive immune response for different infections. The CTL-generated cellular immune response eliminated the infected cells and also the antigens circulating in body (23). On the contrary, HTL epitopes are required to generate both cell-mediated and humoral immune responses (24). They also develop the memory HTLs that activated CTLs and antibody-producing B cells (25). Thus, the evaluation of HTL epitopes (MHC II) and CTL (MHC I) epitopes of the multi-epitope vaccine construct is pre-requisite to design an efficient vaccine candidate capable of elucidating an effecting cell-mediated and humoral immune response. We used the online IEDB server for predicting the MHC-I & MHC-II epitope(s).

#### 3.1.1 Prediction of MHC-I epitopes

For MHC-I, the prediction of the epitopes of HEV capsid protein, ORF3 and polyprotein was done. The epitopes with < 0.1, 1 and 0.05 percentile score were selected for capsid protein, ORF3 and polyprotein respectively. 93, 4 and 80 epitopes having the percentile score <0.1, 1 and 0.05 percentile score in HEV capsid protein, ORF3 and polyprotein respectively were selected and used for further analysis. These epitopes were further evaluated for their immunogenicity and antigenicity. After analysis 3, 4 and 5 epitopes have antigenicity score of more than 0.4 and are projected to be probable antigens for HEV capsid protein, ORF3 and polyprotein **(Table 1)**.

**Table 1:**
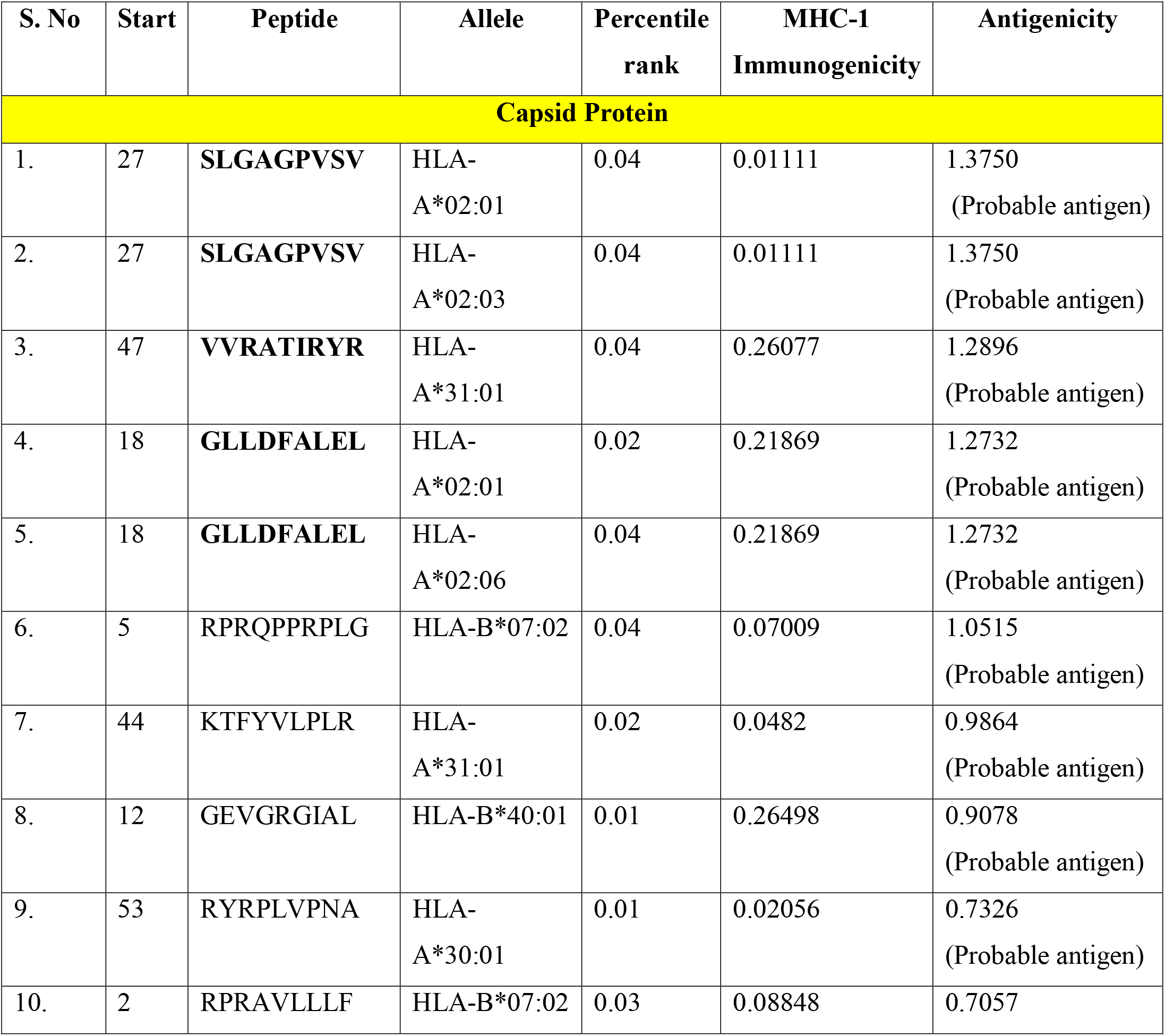
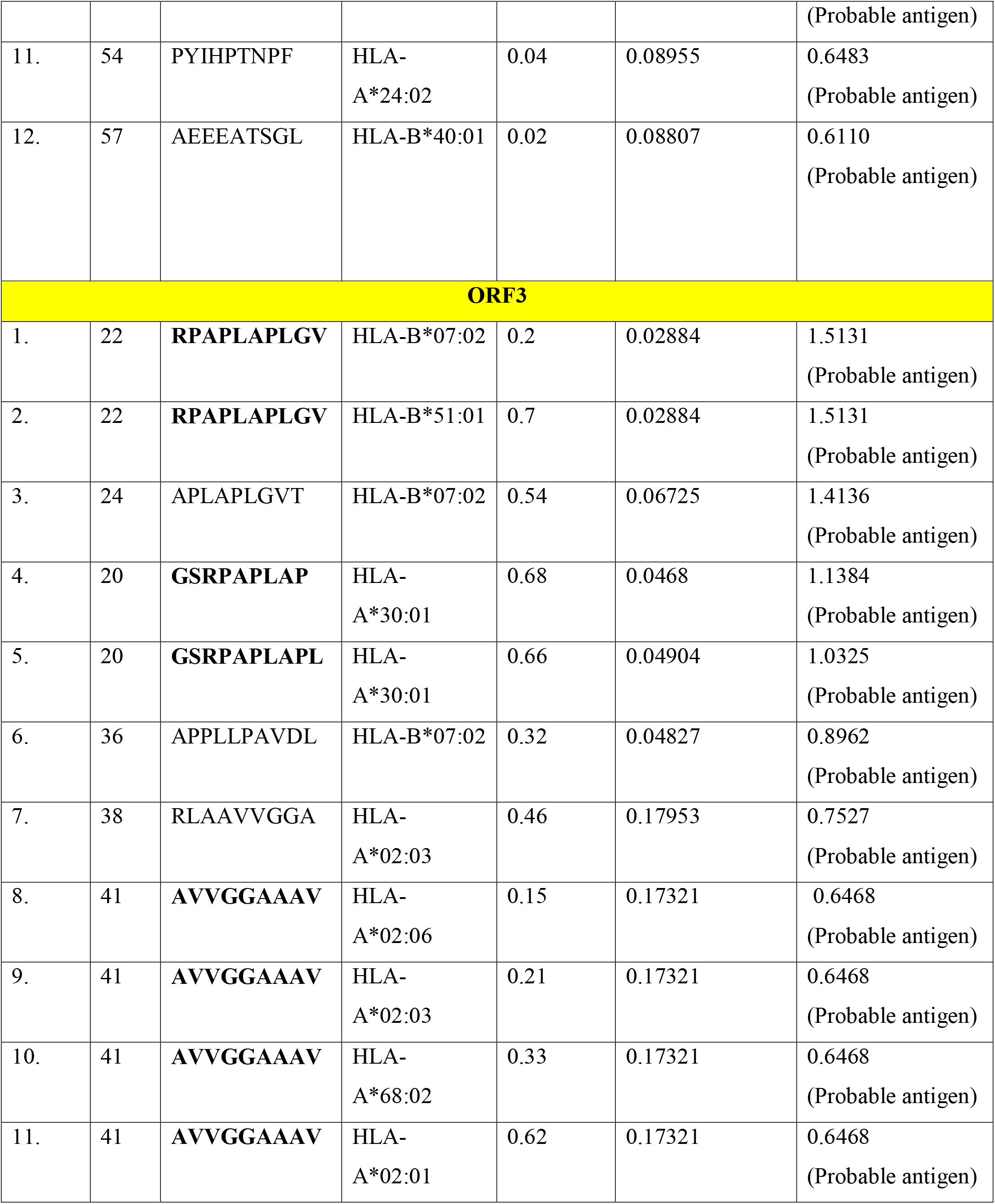
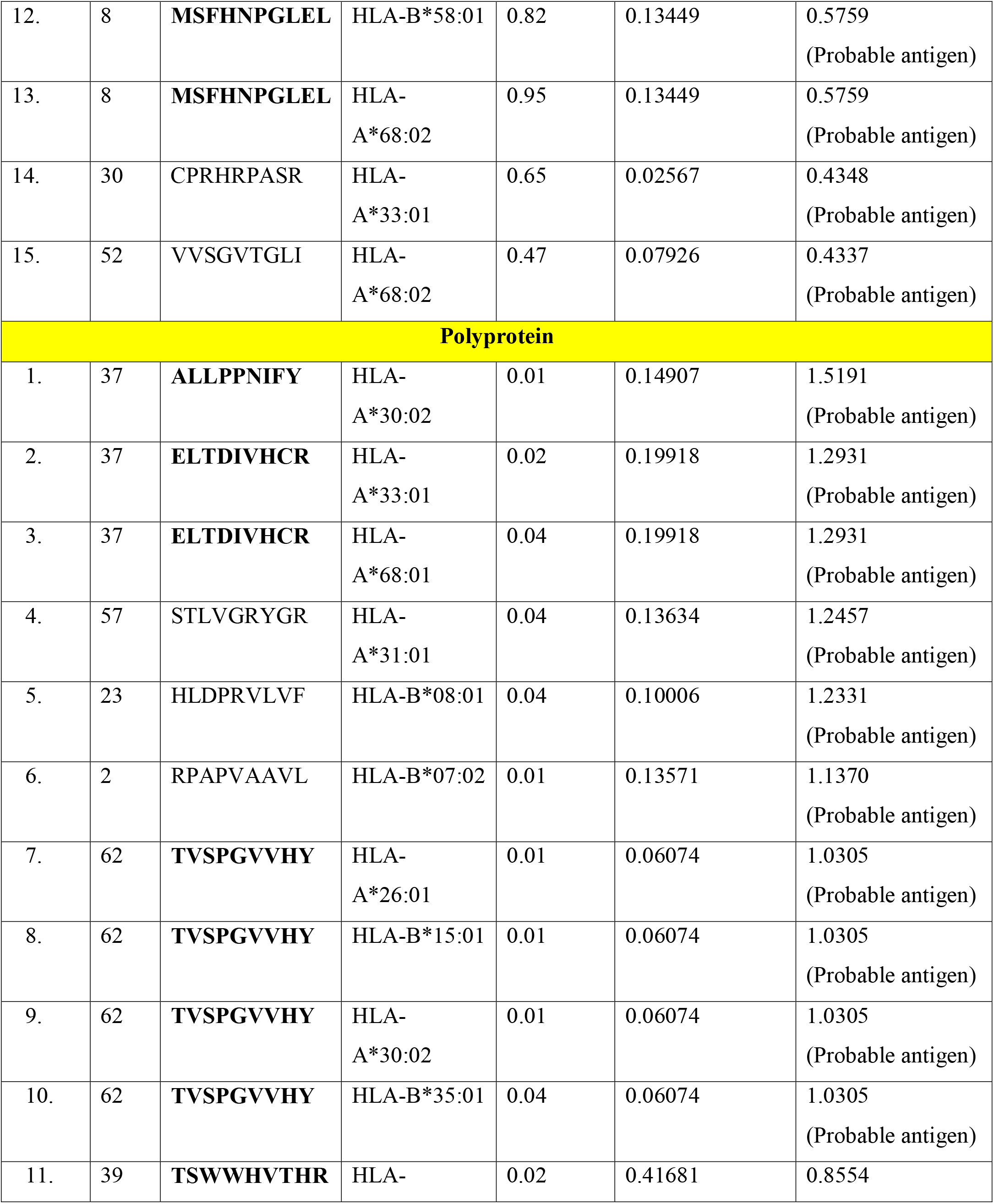
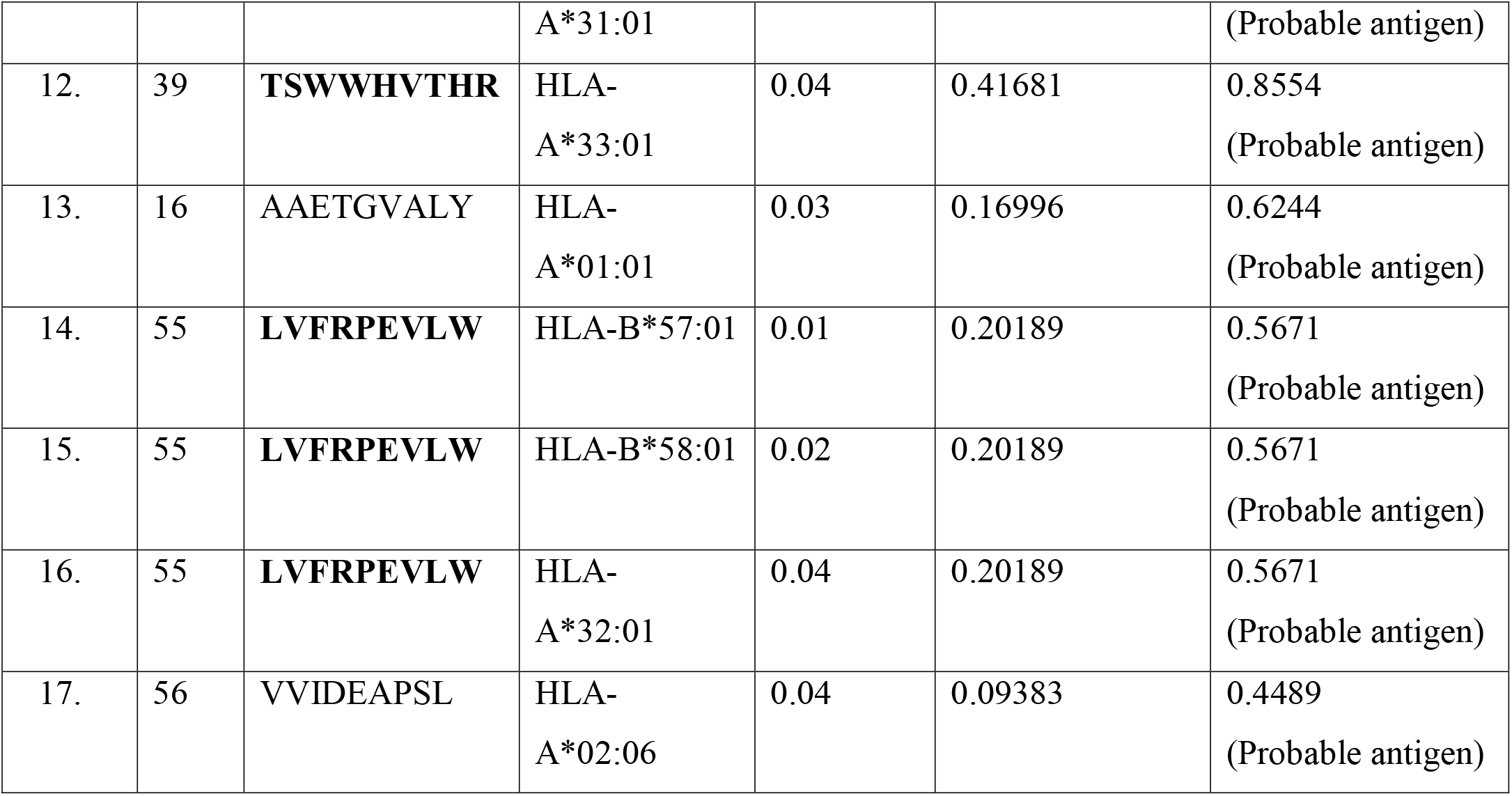
MHC I Epitopes of HEV capsid protein, ORF3 and polyprotein

#### 3.1.2 Prediction of MHC-II epitopes

For predicting the MHC-II epitopes of viral capsid protein, polyprotein and ORF3 7 allele set of HLAs was selected. Epitopes having a percentile score <1 for capsid protein and polyprotein and <0.5 for ORF3 were shortlisted. A total of 9 epitopes from ORF3, 94 from the capsid protein and 157 epitopes were predicted from the viral polyprotein and used for further analysis. The viral epitopes were further evaluated for the positive IFN-γ Score (**Table 2)**. 17 epitopes from capsid protein, 9 from ORF3 and 9 from polyprotein had positive IFN-γ scores **(Table 2)**.

**Table 2:**
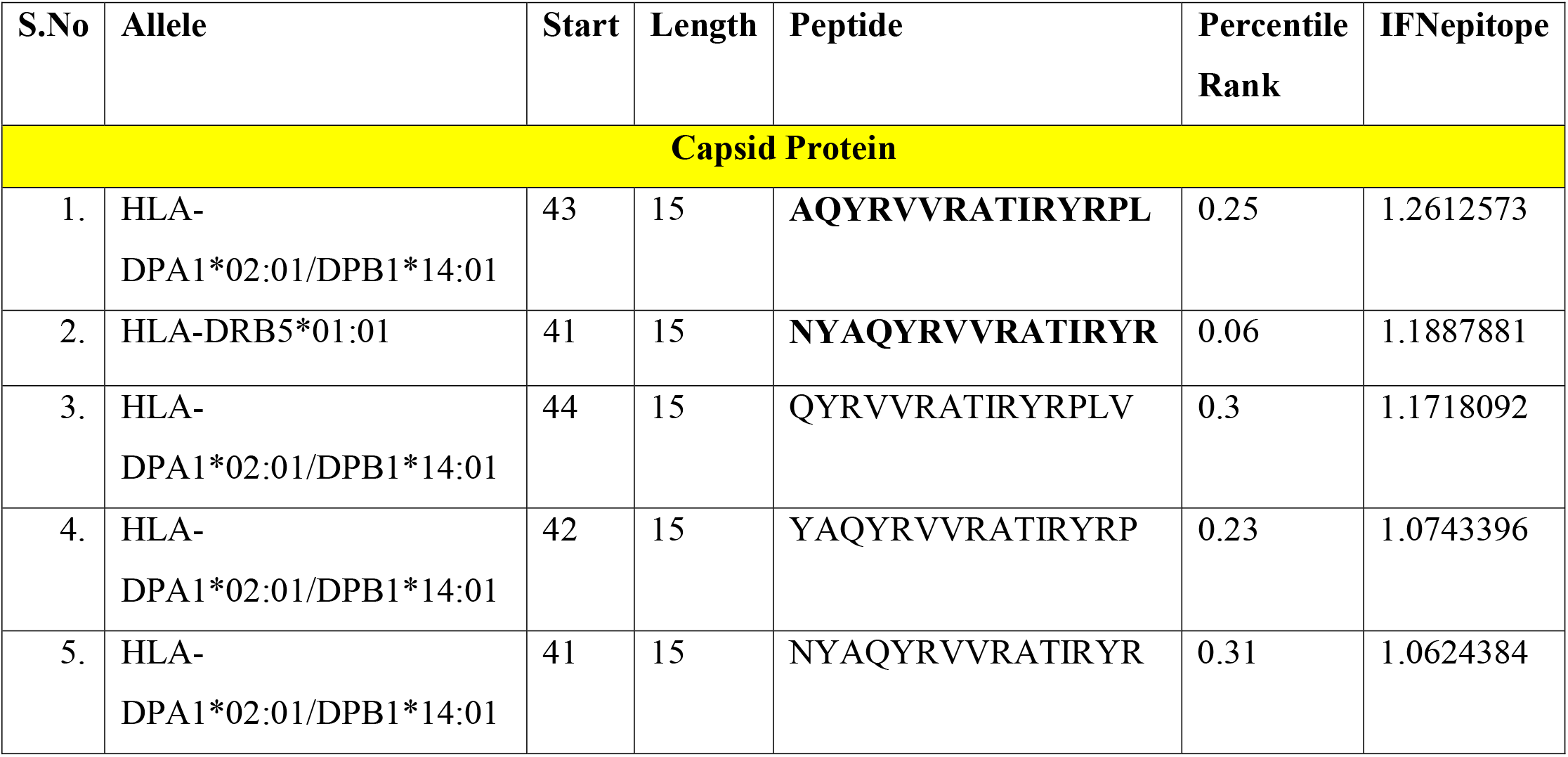
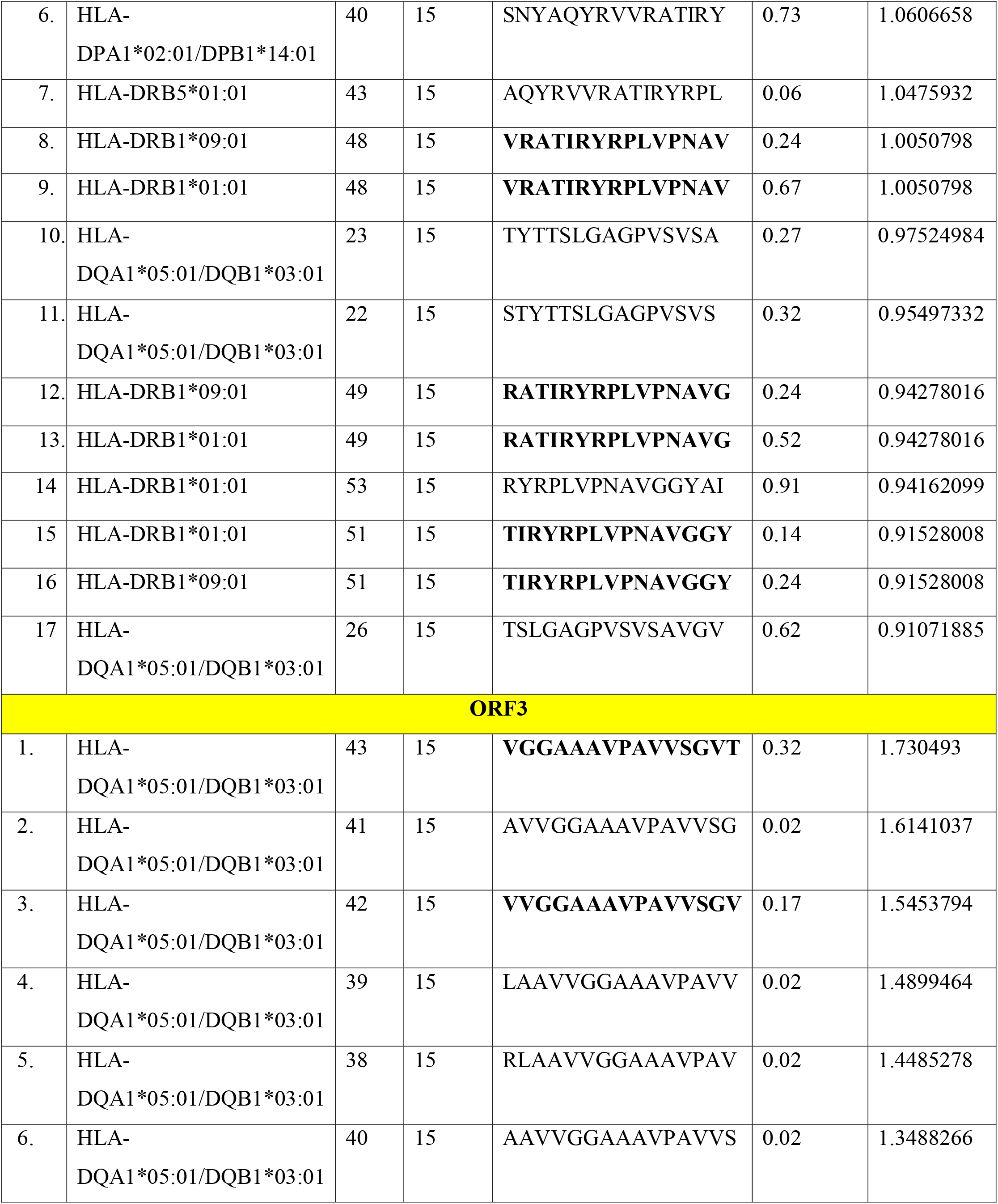
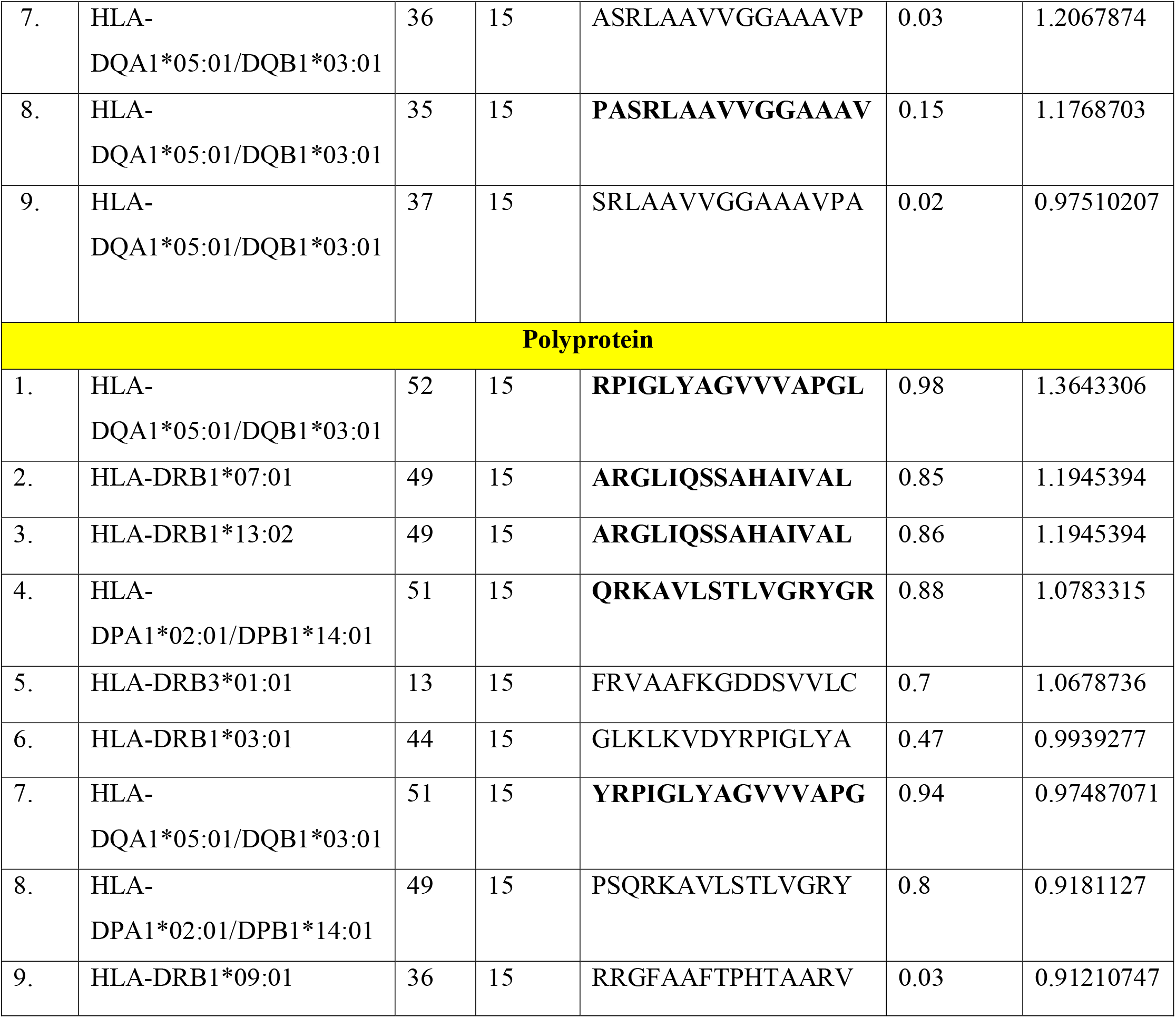
MHC II Epitopes of capsid protein, ORF3 and polyprotein of HEV.

### 3.2 Population coverage analysis

The predicted vaccine construct was expected to cover a wide human population across the world. The IEDB population coverage tool (http://tools.iedb.org/population/) was used to evaluate the distribution of MHC I and MHC II epitope alleles used in the construction of the vaccine construct. The MHC I and MHC II epitopes used for population coverage analysis can cover 86.85% of the world population **(Figure 3A and 3B)**. The population coverage percentage in Central Africa, East Africa, South Africa, North Africa, and West Africa were 62.62%, 71.8%, 66.5%, 77.2% and 77% respectively. In East Asia, Northeast Asia, South Asia, Southeast Asia and Southwest Asia the percentage was 80.8%, 63.82%, 71.46%, 55.76% and 71.48%. In Europe the percentage was 84.74%, in West Indies, it was 81.52%, in Central America, North America and South America it was 4.14%, 83.87% and 65% respectively **(Figure 3C)**. In Oceania, this percentage was 34.28%. Altogether these results indicate the predicted epitopes cover a wide range of the human population and are suitable to be used as a vaccine construct.

### 3.3 Construction and characterization of the multi-epitope vaccine

The competent epitopes of ORF3, capsid protein and polyprotein of HEV were selected and connected with precise linkers for MHC class I, MHC class II epitopes and the adjuvant. The 45 amino acid long adjuvant β-defensin was linked to the N terminal of the MHC-I alleles with EAAAK linker. The MHC-I epitopes of ORF3, capsid protein and polyprotein of HEV were linked via AAY linker while the MHC-II epitopes were linked via GPGPG linker creating the vaccine construct **(Figure 2)**.

**Figure 1:**
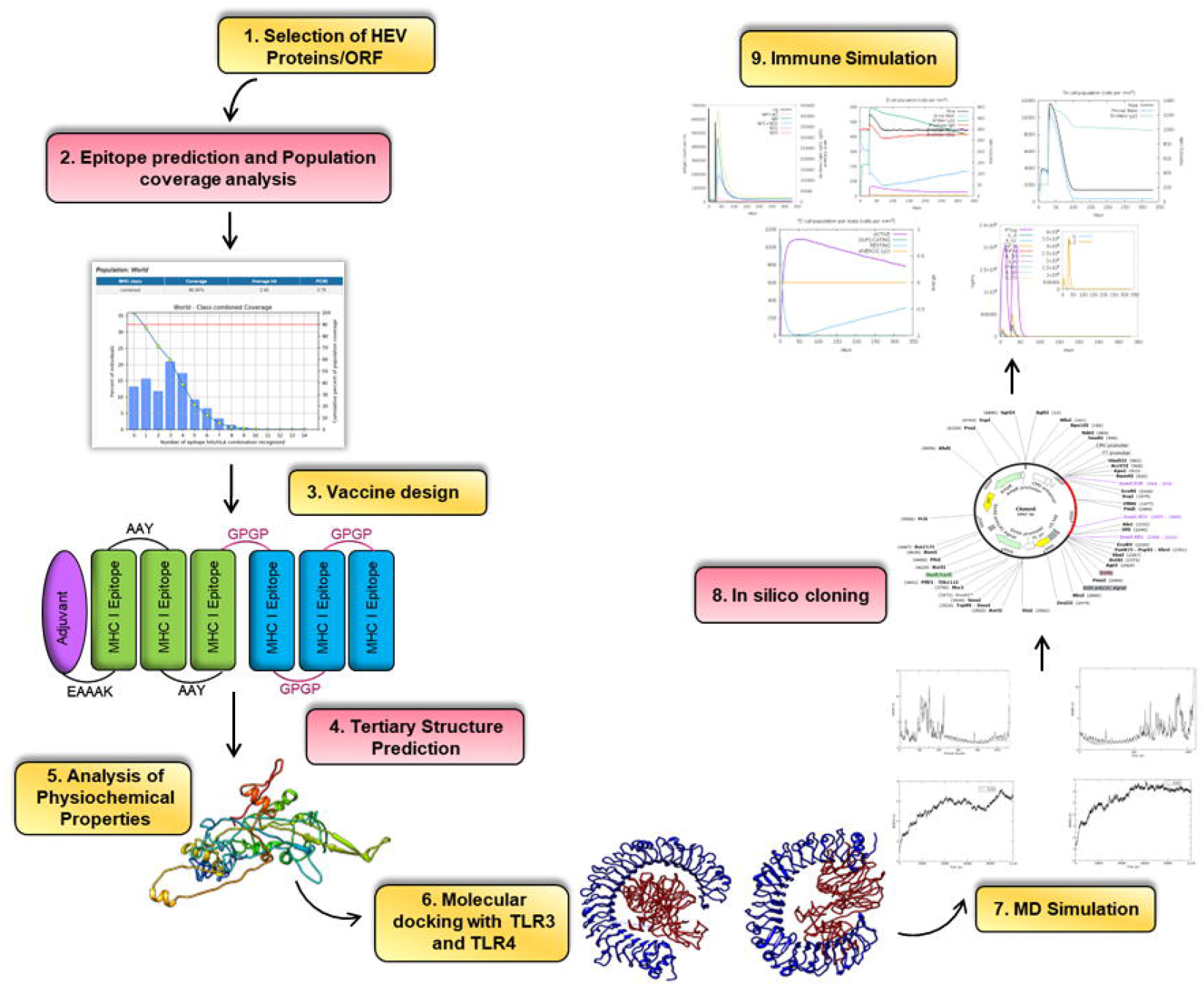
An overview of multi-epitope vaccine construct designing using different *in-silico* tools.

**Figure 2:**
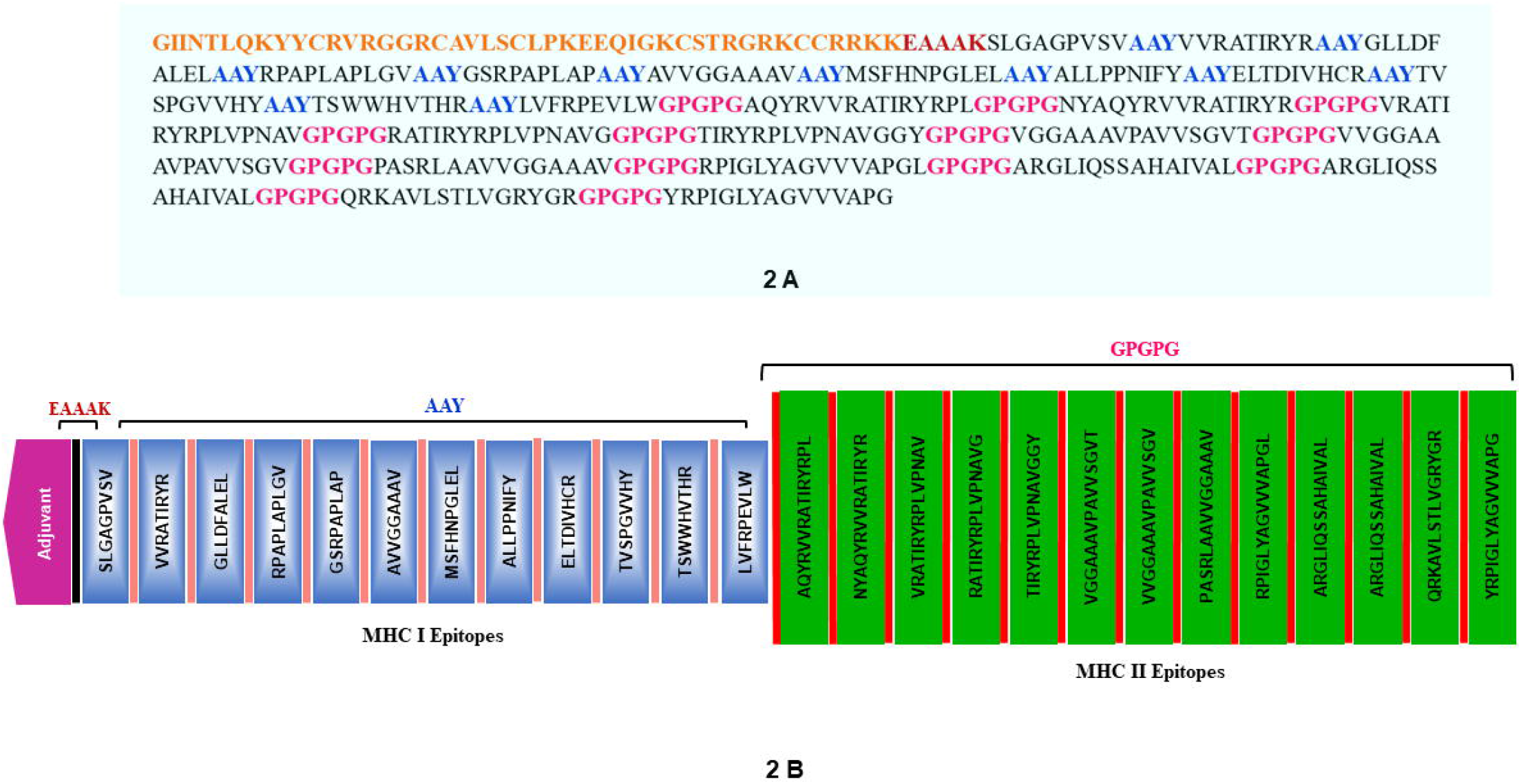
The prepared multi-epitope vaccine construct depicting the amino acid sequence and MHC-I and MHC-II epitopes joined via respective linkers and adjuvant.

### 3.4 Prediction of Linear B cell epitopes

The B cell epitopes of the vaccine construct were predicted by the ABCpred server with 0.8 cut-off score. A total of 29 potential B cell epitopes were identified in the multi-epitope vaccine construct **(Table 3)**. The B cell epitopes with the highest score were GGPGPGTIRYRPLVPN, AYTSWWHVTHRAAYLV and GGYGPGPGVGGAAAVP with scores of 0.93, 0.93, 0.91 respectively.

**Table 3:**
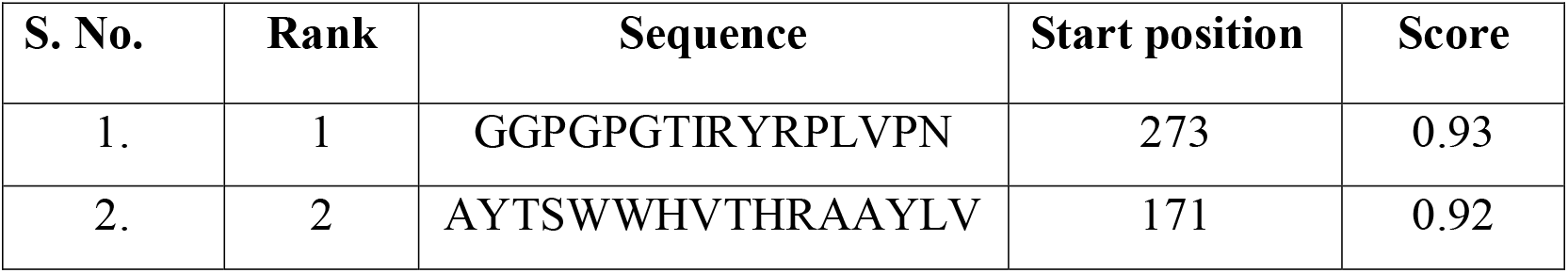
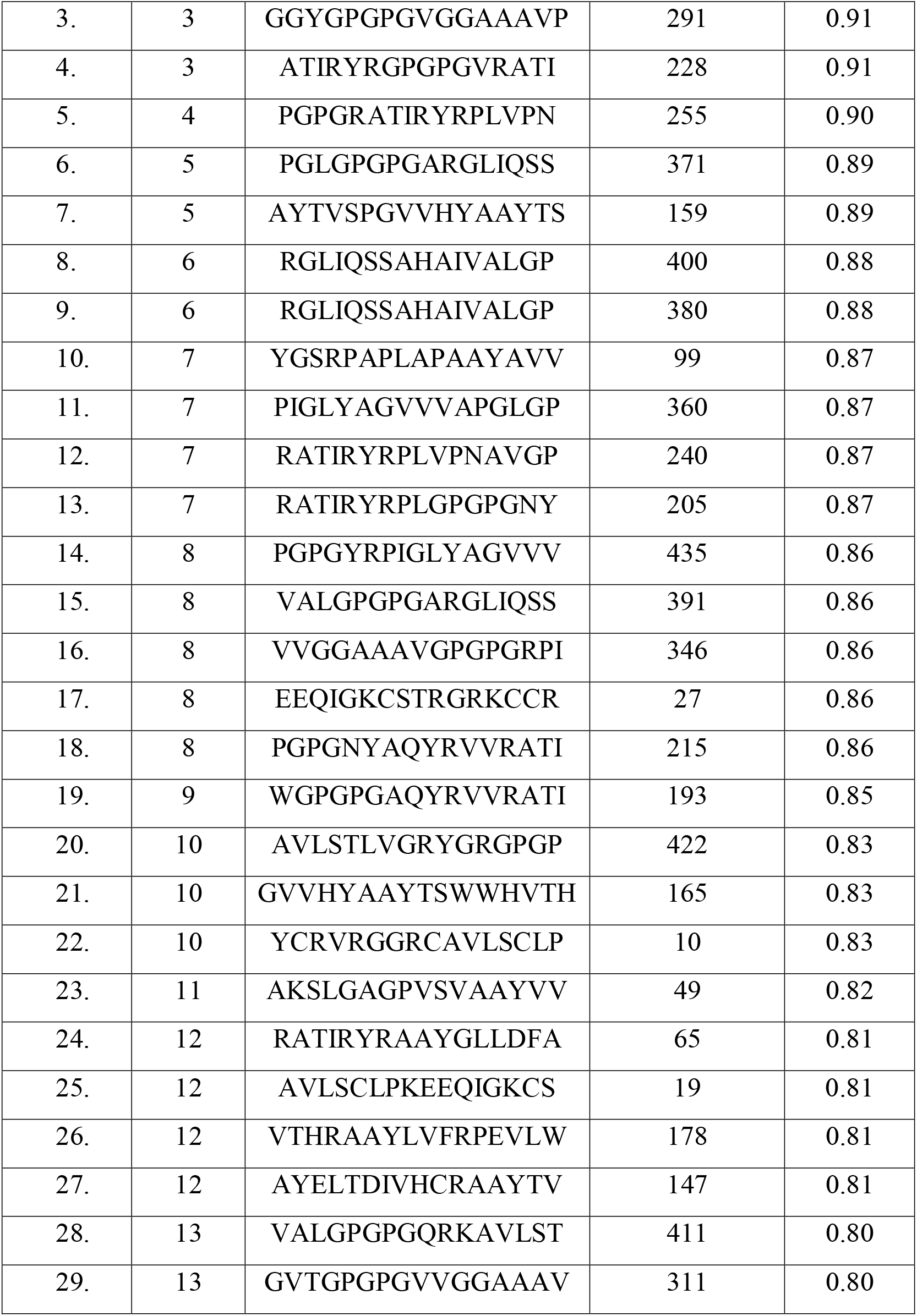
B cell epitopes of the multi-epitope vaccine construct

### 3.5 The analysis of different physiochemical properties of the multi-epitope vaccine construct

The ProtParam analysis of different physiochemical properties of the proposed construct gave favorable results. The 453 amino acid long vaccine construct has a molecular weight of about 45794.23 g/mol and a theoretical pI of 10.5 depicting its basic nature. The proposed construct resulted to be thermostable with an aliphatic index of 91.81 and an average half-life in 30 hours *in vivo* (in mammalian reticulocytes), whereas, >10 & >20 hours in *E. coli* & yeast respectively **(Supplementary data)**.

### 3.6 Allergenicity and Antigenicity

Using the AllerTOPv2.0 algorithm, the multiepitope vaccine construct was predicted to have nonallergenic properties. The Vaxijen v2.0 projected antigenicity score of our multiepitope construct was 0.6327 with >0.4% of threshold value indicating the antigenic properties of the construct.

### 3.7 The *in-silico* vaccine construct favored the secondary and tertiary structure

Upon analysis of the secondary structure for the long 453 amino acid multi-epitope vaccine, it was determined that out of the 453 amino acids, 136 (30.02%) formed alpha helix, 112 (24.73%) beta strands and coil were formed by 205 (42.25%) **(Figure 4)**. A total of 5 models were predicted by I-TASSER using 10 threading templates with Z-scores and models were selected based on their high confidence C values, which have a range between -1.71 and 3.86. The selected models were analyzed for the Ramachandran plots and found that 91.1% of residues were located in the most favored regions and additional allowed regions.

**Figure 3:**
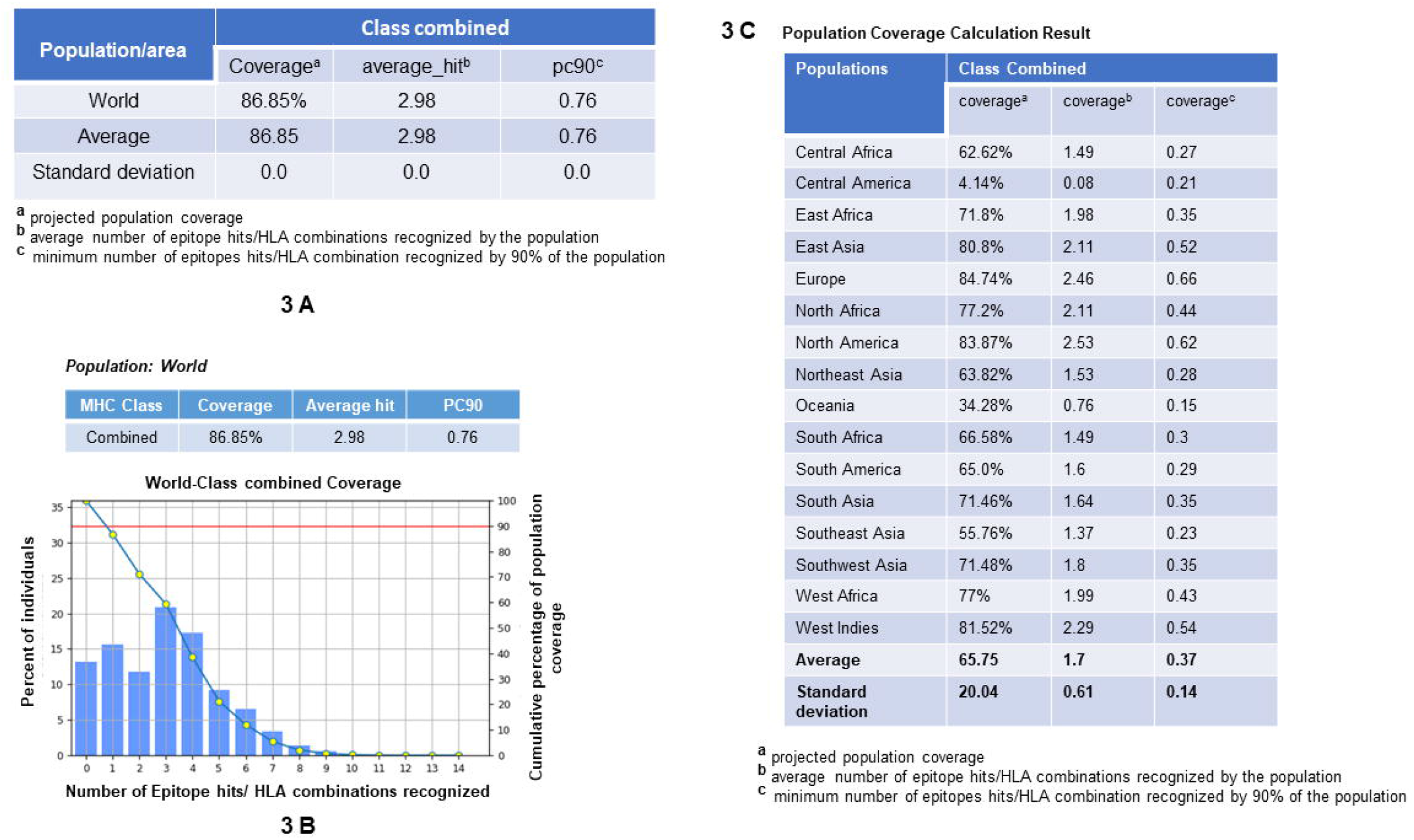
Population coverage analysis of the multi epitope vaccine construct

**Figure 4:**
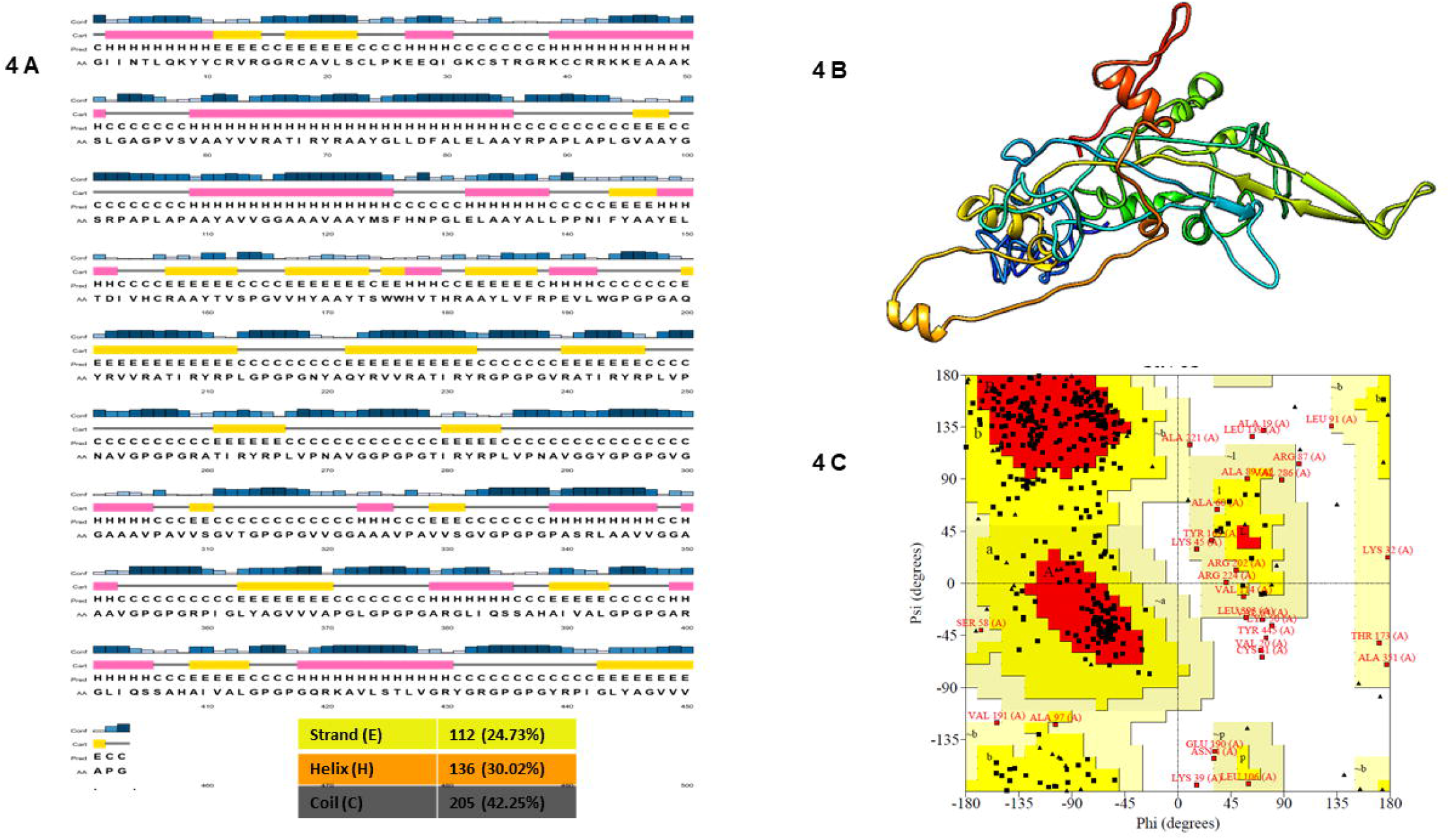
Predicted 2D and 3D structure of the *in-silico* construct (3a. Projected 2D structure of the proposed vaccine construct, 3b. Projected 3D structure of the proposed vaccine construct, 3c. Ramachandran Plot examination of the proposed vaccine construct).

### 3.8 TLR3 and TLR4 interaction with multi-epitope vaccine construct

The *in-silico* vaccine construct with the nominated epitopes against the HEV was docked against TLR4 (3FXI) and TLR3 (2A0Z). The docking results displayed a steady interface between the projected vaccine construct and the TLR3 and 4. These interactions were envisioned in LigPlot **(Figure 5)**.

**Figure 5:**
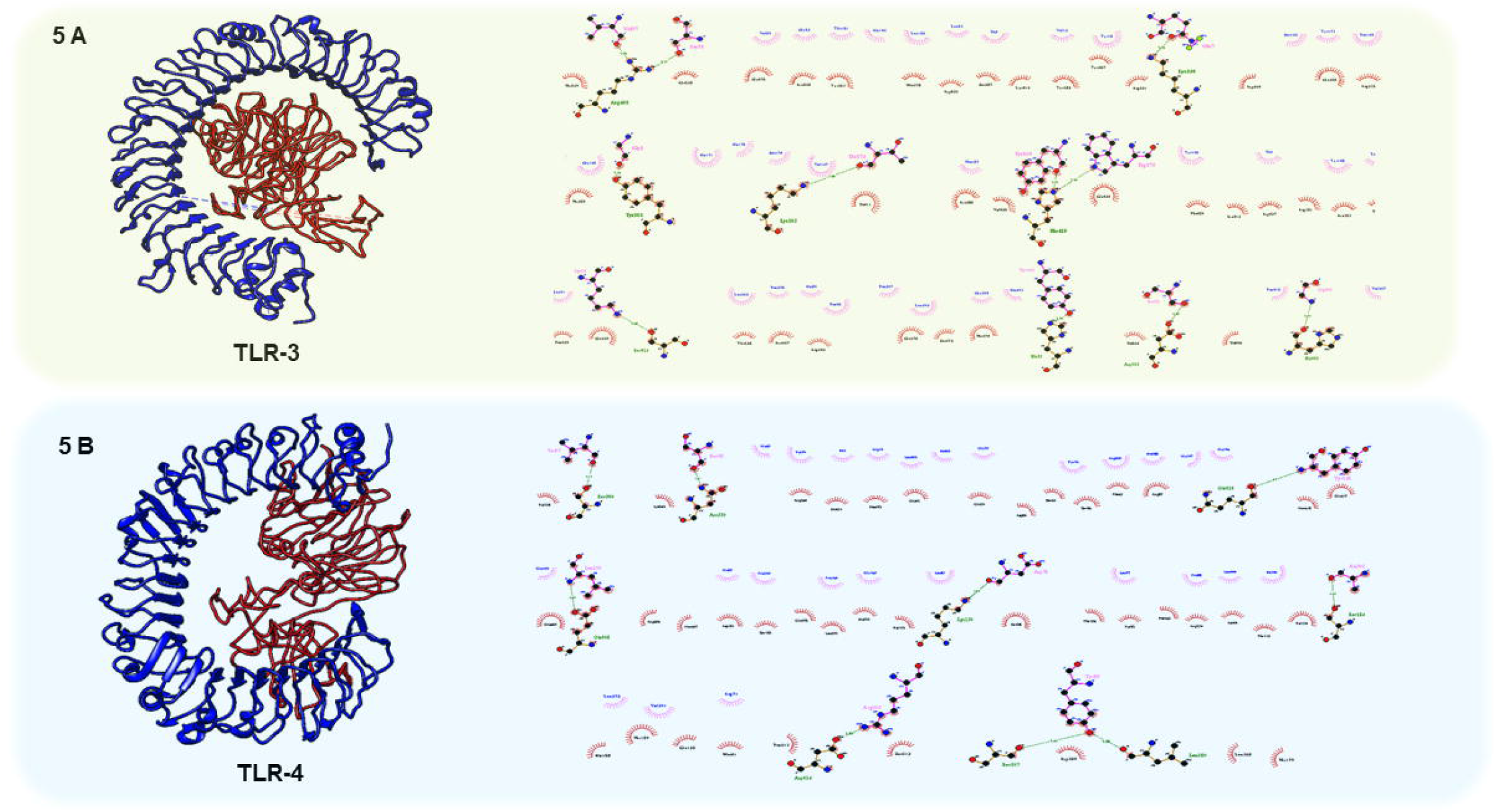
Molecular docking results of the multi-epitope vaccine construct with TLR 3 & TLR 4 receptors. (**4A**. Representation of the docking interaction and conformation of the TLR 3 & multi-epitope vaccine construct in Ligplot analysis, **4B**. Representation of the docked interaction and conformation of the TLR 4 & multi-epitope vaccine construct in Ligplot analysis).

### 3.9 MD simulation results support a steady ligand-receptor interaction

The molecular dynamics analysis revealed the interaction dynamics and insights into the flexibility of the binding site. The stability of the simulations was assessed by root mean square deviation (RMSD). The RMSD indicates the changes in the structure during the MD compared to a reference point usually the initial point. It is an important indicator of the stability of a simulation. In the current study, RMSD plots for the generated complexes showed that each complex got stabilized quickly. The RMSD remained stable throughout the simulation as indicated by a narrow movement of the RMSD curve within 2Å. The root mean square fluctuation (RMSF) was used to check the flexibility of the individual residues (and in turn the secondary structural elements). The residues making strong interactions are usually show less flexibility as compared to other amino acid residues. Here, it can be seen from RMSF plots **(Figure 6A and 6B)** that the beta sheets of both TLR3 and TLR4 are showing very less fluctuations (in a wavy pattern) due to strong hydrogen bonding. The HEV though is showing greater fluctuations due to the presence of many loops. A comparison of the RMSD plots of TLR3 and TLR4 shows that the TLR3 complex has lower RMSF i.e. comparatively lesser fluctuations. It indicates better binding among the interaction partners and a comparatively stronger complex.

**Figure 6:**
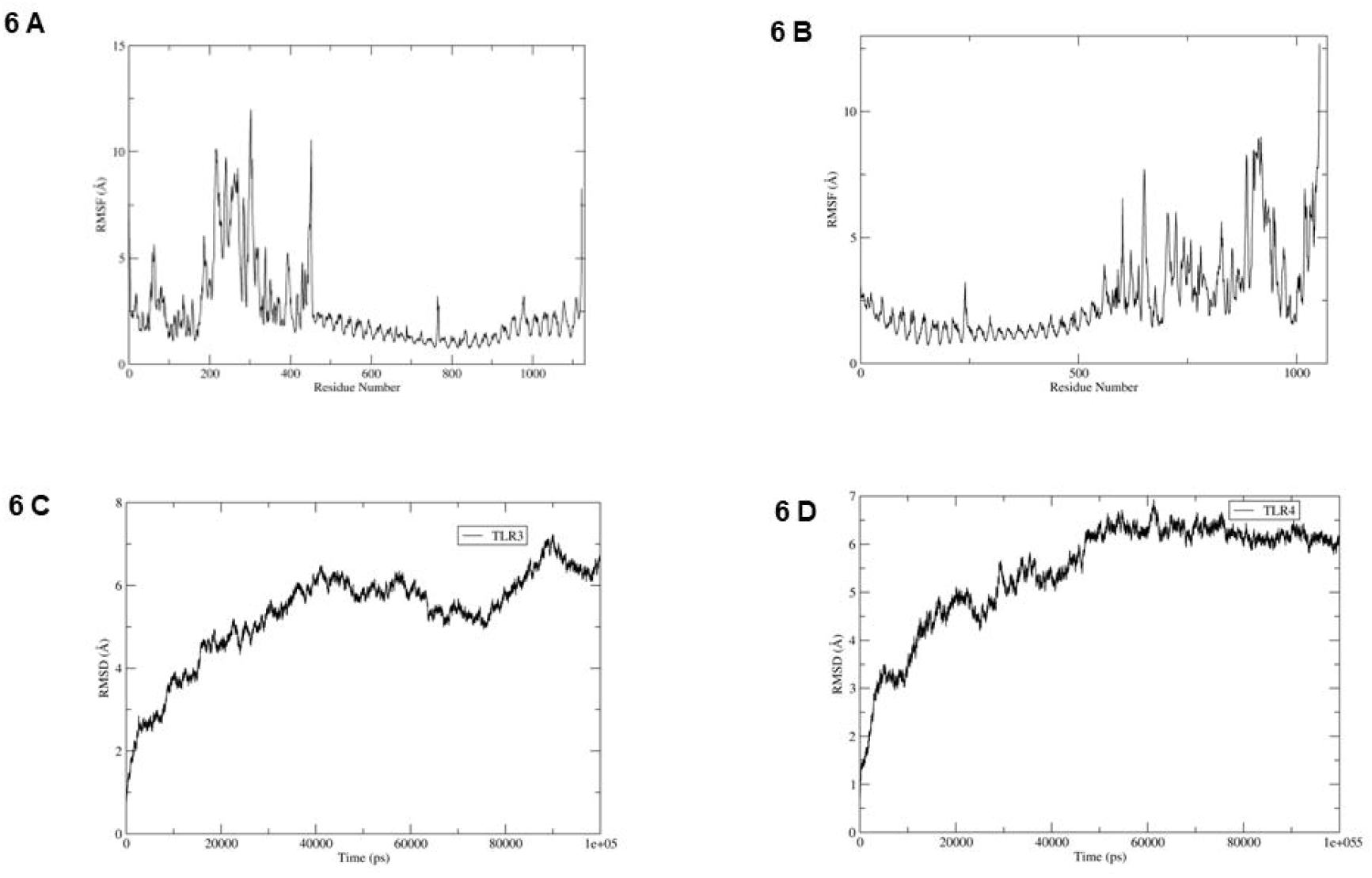
MD Simulation Results (**5A**. The RMSF curve for TLR3-HEV construct, **5B**. RMSF curve for TLR4-HEV construct. The beta sheets of the TLR3 and TLR4 show limited flexibility. The flexibility of the interface binding residues is also less indicating stable interactions, **5C**. RMSD curve for TLR3-HEV construct, **5D**. RMSD curve for TLR4-HEV construct). The figures clearly depict that the complexes got stabilized quickly.

The salt-bridge analysis in the construct-TLR3 complex showed residues at the binding interface have strong interactions especially GLU570_TLR3-LYS50_construct and ASP592_TLR3-LYS50_construct with occupancy of 67% and 43% respectively **(Figure 7A)**. On the other hand, the binding, interactions were less strong in the construct-TLR4 complex as compared with the construct-TLR3 complex. The GLU603_TLR4-LYS26_construct, and ASP78_construct-LYS130_TLR3 interactions showed 28% and 24% occupancy respectively. The salt-bridge interactions are shown in **Figure 7B**.

**Figure 7:**
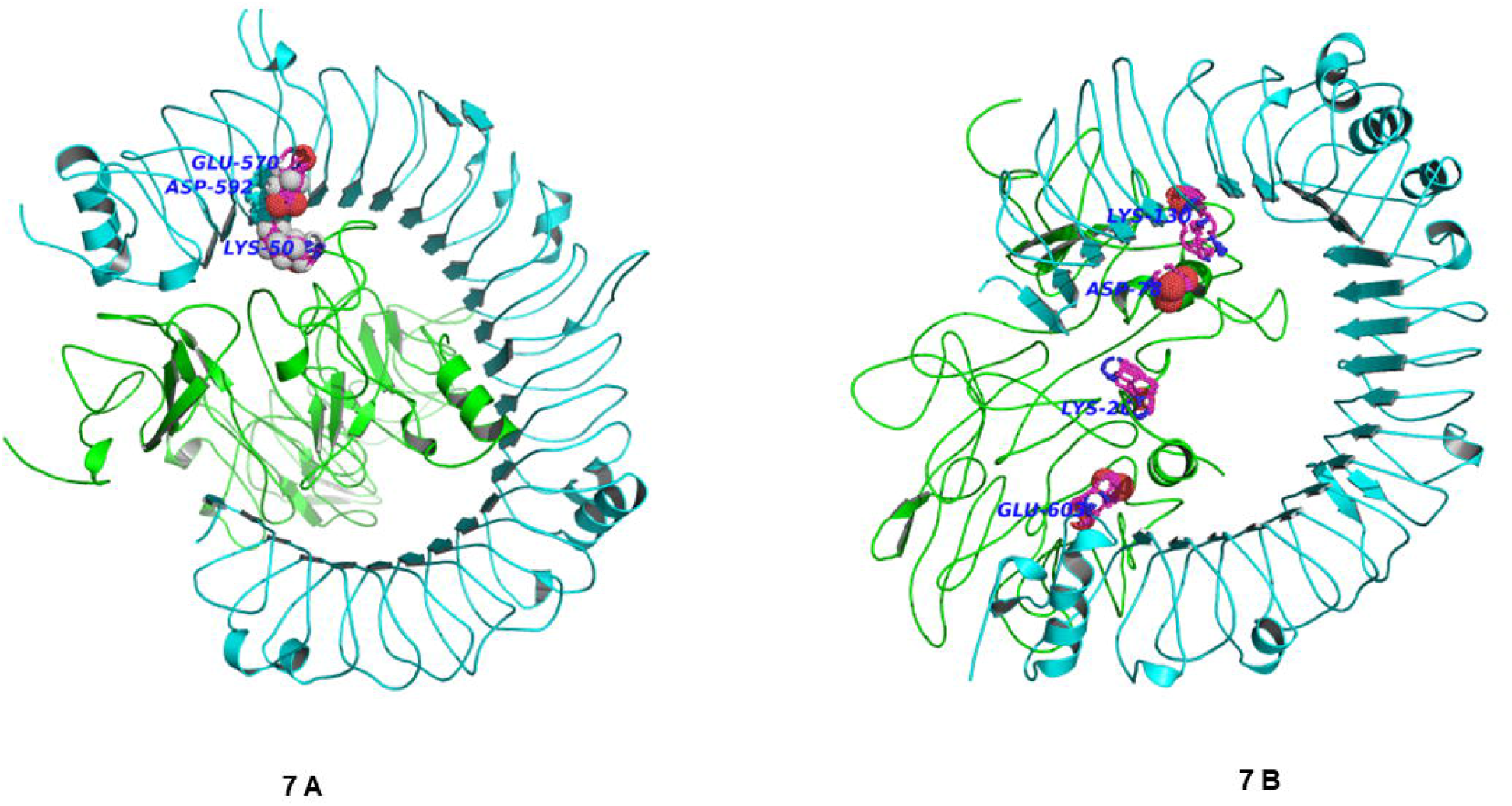
Salt bridges between vaccine construct and TLR3 and TLR4 (**7A**. The complex of TLR3-vaccine construct, **7B**. The complex of TLR4-vaccine construct. The three most stable salt bridges are shown for both of the complexes. The salt bridges anchor the two proteins through stable salt-bridge interactions.

### 3.10 Cloning of the vaccine construct in pcDNA3.1/V5/His-Topo vector through *in-silico* tool

*In-silico* vaccine candidate was codon optimized using the Jcat server for higher protein expression in *E.coli*. The vaccine had 1359 nucleotides and 79.47% GC content. The construct was cloned into the pcDNA3.1/V5/His-Topo expression vector by using the SnapGene software **(Figure 8)**.

**Figure 8:**
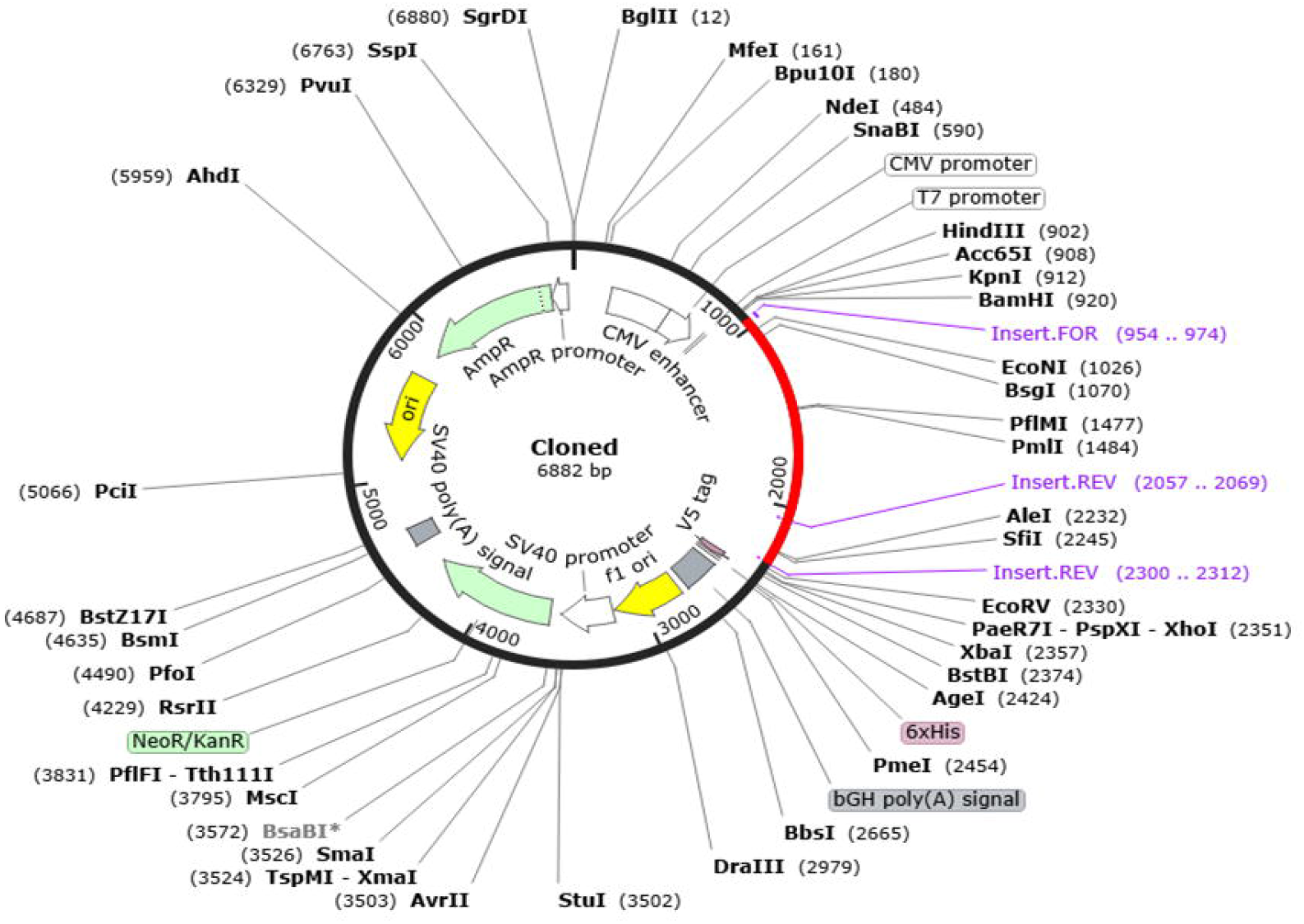
*In-silico* cloning of multi-epitope vaccine in pcDNA3.1/V5/His-TOPO/LacZ vector

### 3.11 Immune simulation results indicate an effective antibody and cytokine response

The immune simulation results from the C-ImmSim displayed an efficient immune response triggered by the *in-silico* vaccine construct. These results were indicated by upregulated IgM level and IgG1+IgG2, IgM and IgG+ IgM expression. In addition, the upregulated cytotoxic T cells and Helper T cells results also confirmed an appropriate immune response triggered by the multi-epitope vaccine construct. Also, the cytokine production and IFN-γ along with B and T cells were observed **(Figure 9)**.

**Figure 9:**
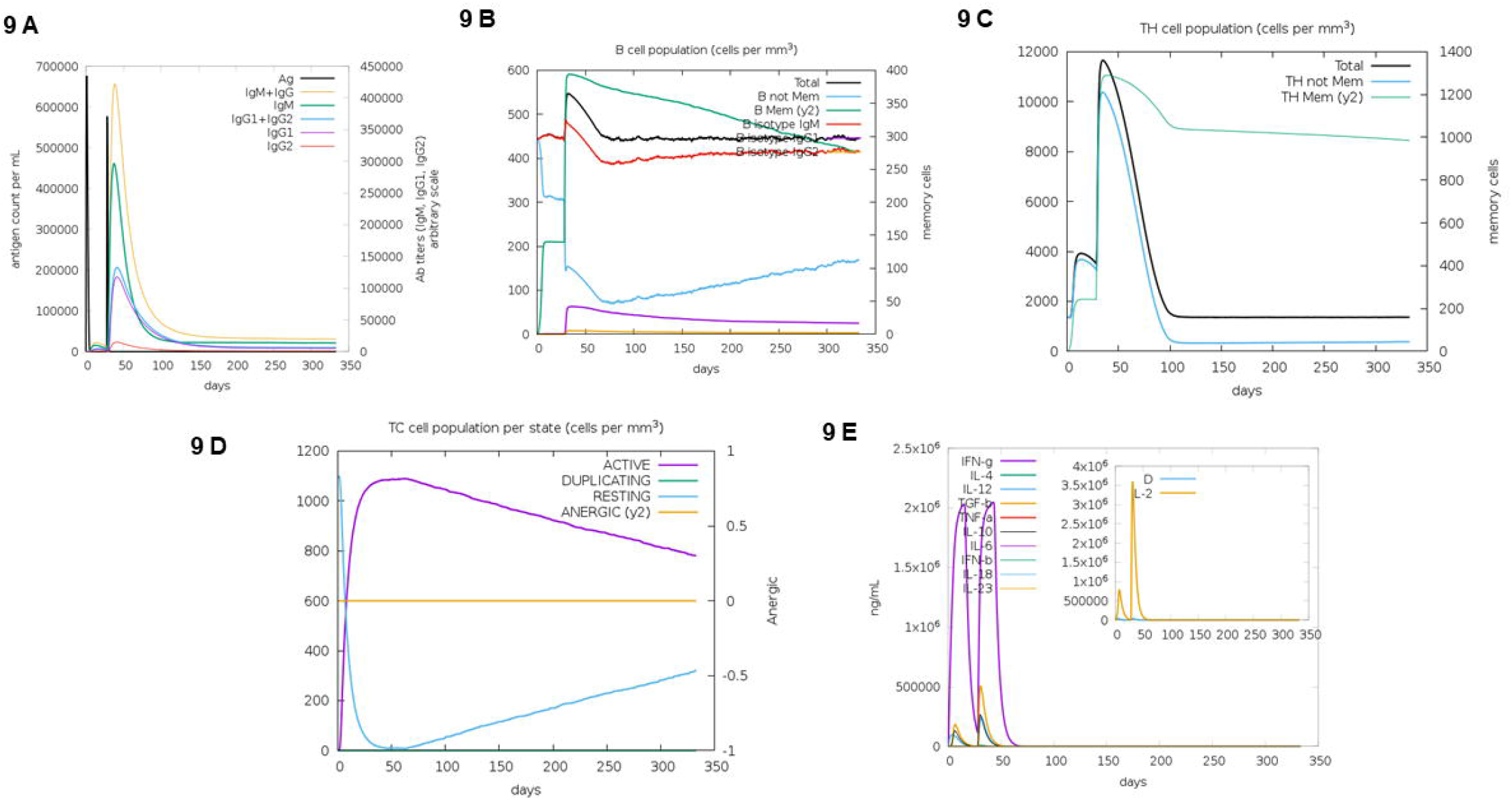
Immune stimulation results of the proposed vaccine construct generated through C-ImmSim server after 3 dose injections of vaccine construct (**7A**. Generation of the immunoglobulin after antigen inoculations (black vertical lines); detailed subclasses areshownd as colour peaks, **7B**. Evolution of B-cell populations, **7C**. Th cell population evolution, **7D**. Populations of the Cytotoxic-T cells, **7E**. Cytokines & interleukins production).

## 4. Discussion

HEV infection is associated with about 50% of hepatitis cases in Asian countries like India and China (26). The acute hepatitis E symptoms are indistinguishable from other acute viral hepatitis, including anorexia, mild, nausea, fever and vomiting. Other symptoms include skin rash, abdominal pain, or joint pain and the patients can even advance jaundice with hepatomegaly and dark urine. In infrequent instances, fulminant hepatitis can also occur (27). The rate of fulminant hepatitis significantly increases in HEV-infected second or third trimester pregnant women resulting in fetal loss and acute liver failure with a mortality rate as high as 25%. Recently, chronic HEV infections are becoming a substantial clinical manifestation in immunosuppressed individuals, especially in organ transplant patients (28), which could allow the virus to mutate and potentially enhance its transmissibility and pathogenicity (29). After more than three decades of HEV discovery and genome characterization, our understanding of HEV pathogenesis and prevention is still restricted. Furthermore, HEV-associated complications are no longer limited to developing countries and is now a concern in developed countries as well (30).

Hecolin® is the lone vaccine available against HEV that is only being administered in China. It is a prophylactic vaccine based on HEV genotype 1 capsid protein. Though, it is not clear if it will protect humans or domestic animals from zoonotic HEVs. In addition, quasi-enveloped HEV particles explain why only capsids targeting vaccines have not worked (30).

Even though ongoing research continues to reveal new insights about HEV pathogenesis, we still lack proper therapeutical options against it. A major reason for the unavailability of effective live attenuated and/or killed vaccines is the absence of a permissive HEV cell culture system (31). Thus, the major emphasis of HEV vaccine development is on molecular approaches, mainly expressing the viral proteins in various expression systems (32). This multi-epitope design is a new approach that can counter current limitations in HEV vaccines, but it needs further investigation.

The conventional vaccine development approach is an expensive and time-consuming process requiring culturing of the pathogen in the laboratory (33). Reverse vaccinology exploits genomic information and computational analysis for vaccine development without culturing the pathogen (34). Prediction of the T and B cell epitopes, antigen processing analysis, antigenicity study, allergenicity analysis, toxicity prediction, and TLR-peptide docking are crucial steps of vaccine designing against several viruses (35).

The purpose of the current study was to develop a multi-epitope vaccine against HEV using a reverse vaccinology approach. The current study uses highly immunogenic and antigenic MHC I and MHC II peptides from the capsid protein, ORF3 and polyprotein of the HEV virus. The three selected targets are crucial for viral survival and replication. The capsid protein is needed for viral packaging, and the ORF3 translation product facilitates viral immune evasion. Furthermore, the HEV polyprotein codes for several crucial non-structural proteins and altogether targeting these viral components can be useful to limit HEV infection. We used the highly immunogenic epitopes of all three targeted proteins with a percentile score <1 for capsid protein and polyprotein and <0.5 for ORF3. These viral epitopes were further evaluated for the positive IFN-γ Score and 17 epitopes from capsid protein, 9 from ORF3 and 9 from polyprotein had positive IFN-γ score. These epitopes were used to construct the multiepitope vaccine using appropriate linkers and adjuvant. The proposed vaccine construct was analyzed for B-cell epitopes via the ABCpred server. GGPGPGTIRYRPLVPN, AYTSWWHVTHRAAYLV, GGYGPGPGVGGAAAVP were the most potent epitopes with 0.92, 0.91 and 0.9 scores respectively. The complete *in-silico* construct had a chemical formula of C_20_^79^H_3291_N_609_O_545_S_8_ and a molecular weight of 45.79kDa. The construct exhibited a theoretical PI of 10.5 implying the basic nature of our construct. In addition, the Grand average of hydropathicity (GRAVY) of the construct was 0.206, the aliphatic index was 91.81 and the instability index (II) is computed to be 29.72 that classifies our construct as stable. The secondary and tertiary structure analysis of the multi-epitope vaccine construct supports the stability of our construct. The 453 amino acid long multi-epitope vaccine has136 (30.02%) alpha helix forming amino acids, 112 (24.73%) beta strands forming amino acids and 205 (42.25%) coil forming amino acids. Total 5 different models were predicted by I-TASSER software and selection was based on the high confidence C values. These modes were validated by the Ramachandran plot and 91.1% of residues were located in the most favored and additional allowed regions.

The current multi epitope construct is non-allergic and antigenic signifying its value for the induction of host immune responses without an allergic reaction. Furthermore, the docking results presented the stable interaction between the vaccine construct and the TLR-3 & TLR 4, implying a higher affinity of the construct towards TLR-3 & TLR-4. In addition, our immune simulations resulted supported the efficacy of the vaccine construct by displaying an increased T-cell and B-cell production for several months along with T-Helper cell. The cytokine profile triggered by the *in-silico* construct showed an increased IFN-γ level from first injection onwards. The expression level reached the peak after repeated exposure of the antigen. The construct was codon optimized and cloned into pcDNA3.1/V5/His-Topo *E. coli* expression vector.

Previous studies on anti-HEV vaccine mainly targeted the capsid protein (encoded by ORF2) (36)(37). For the first time we have included HEV ORF3, capsid protein and polyprotein all together for the construction of an efficient, multi-epitope in-silico vaccine construct against HEV. The present vaccine construct is immunogenic, structurally stable, non-allergic and non-toxic. Our proposed vaccine construct had a population coverage percentage of 86.85% implying its wide range of human population coverage. In addition, it was capable of inducing stable and prolonged host immune response. *In vitro* and *in vivo* studies are required to verify the vaccine construct’s safety and effectiveness. If satisfactory results are achieved after the *in vitro* and *in vivo* studies, the proposed vaccine construct might be effective both therapeutically and prophylactically against HEV.

## Acknowledgments

Authors are thankful to the Department of Microbiology, All India Institute of Medical Sciences Bhopal (Madhya Pradesh), India. This work is supported by DBT-Ramalingaswami Re-entry grant BT/RLF/Re-entry/57/2017 to PK

## Author contributions

AK, US, GA and PK conceptualized the manuscript, performed the experiments and wrote the manuscript. GA, PK and AD helped to prepare the manuscript and in the modification of the text. AK, US and PK made the final draft of the manuscript. All authors contributed to the article and approved the submitted version.

## Supplementary data

**Supplementary data 1:** Protparam analysis of the vaccine construct

**Supplementary data 2:** Population coverage analysis of MHC I and MHC II epitopes alleles in different human populations of the world.

