## Supplementary data for "A novel multi-epitope peptide vaccine candidate targeting Hepatitis E virus: an in-silico approach"

**Supplementary data 1: Protparam analysis of the predicted *in silico* vaccine construct**

**Number of amino acids:** 453

**Molecular weight:** 45794.23

**Theoretical pI:** 10.55

**Amino acid composition:**

| | | |
|---|---|---|
| Ala (A) | 74 | 16.3% |
| Arg (R) | 39 | 8.6% |
| Asn (N) | 7 | 1.5% |
| Asp (D) | 2 | 0.4% |
| Cys (C) | 7 | 1.5% |
| Gln (Q) | 7 | 1.5% |
| Glu (E) | 7 | 1.5% |
| Gly (G) | 74 | 16.3% |
| His (H) | 7 | 1.5% |
| Ile (I) | 17 | 3.8% |
| Leu (L) | 32 | 7.1% |
| Lys (K) | 8 | 1.8% |
| Met (M) | 1 | 0.2% |
| Phe (F) | 4 | 0.9% |
| Pro (P) | 53 | 11.7% |
| Ser (S) | 16 | 3.5% |
| Thr (T) | 14 | 3.1% |
| Trp (W) | 3 | 0.7% |
| Tyr (Y) | 29 | 6.4% |
| Val (V) | 52 | 11.5% |

| | | |
|---|---|---|
| Pyl (O) | 0 | 0.0% |
| Sec (U) | 0 | 0.0% |

| | | |
|---|---|---|
| (B) | 0 | 0.0% |
| (Z) | 0 | 0.0% |
| (X) | 0 | 0.0% |

**Total number of negatively charged residues (Asp + Glu):** 9
**Total number of positively charged residues (Arg + Lys):** 47

**Atomic composition:**

| | | |
|---|---|---|
| Carbon | C | 2079 |
| Hydrogen | H | 3291 |
| Nitrogen | N | 609 |
| Oxygen | O | 545 |
| Sulfur | S | 8 |

**Formula:** $C_{2079}H_{3291}N_{609}O_{545}S_{8}$
**Total number of atoms:** 6532

**Extinction coefficients:**

Extinction coefficients are in units of $M^{-1}$ $cm^{-1}$, at 280 nm measured in water.

Ext. coefficient 60085
Abs 0.1% (=1 g/l) 1.312, assuming all pairs of Cys residues form cystines

Ext. coefficient 59710
Abs 0.1% (=1 g/l) 1.304, assuming all Cys residues are reduced

**Estimated half-life:**

The N-terminal of the sequence considered is G (Gly).

The estimated half-life is: 30 hours (mammalian reticulocytes, in vitro).
>20 hours (yeast, in vivo).
>10 hours (Escherichia coli, in vivo).

**Instability index:**

The instability index (II) is computed to be 29.72
This classifies the protein as stable.

**Aliphatic index:** 91.81

**Grand average of hydropathicity (GRAVY):** 0.206

**Supplementary data 2:** Population coverage analysis of MHC I and MHC II epitopes alleles in different human populations of the world.

*Population: Central Africa*

| MHC class | Coverage | Average hit | PC90 |
|---|---|---|---|
| combined | 62.62% | 1.49 | 0.27 |

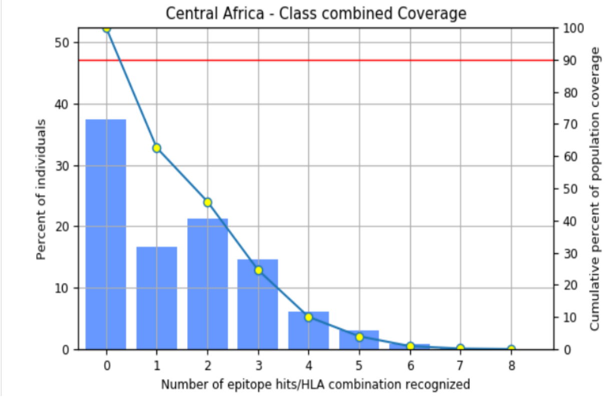


*Population: Central America*

| MHC class | Coverage | Average hit | PC90 |
|---|---|---|---|
| combined | 4.14% | 0.08 | 0.21 |

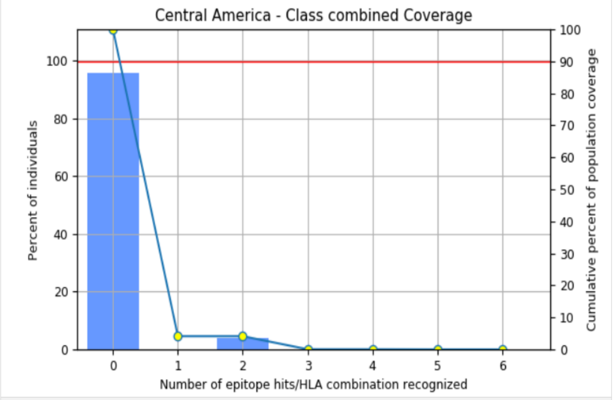


*Population: East Africa*

| MHC class | Coverage | Average hit | PC90 |
|---|---|---|---|
| combined | 71.8% | 1.98 | 0.35 |

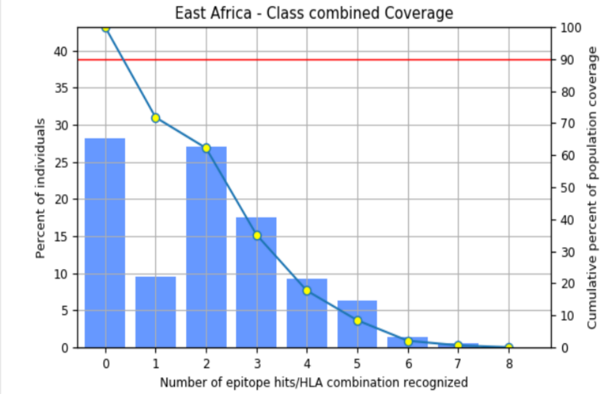


*Population: East Asia*

| MHC class | Coverage | Average hit | PC90 |
|---|---|---|---|
| combined | 80.8% | 2.11 | 0.52 |

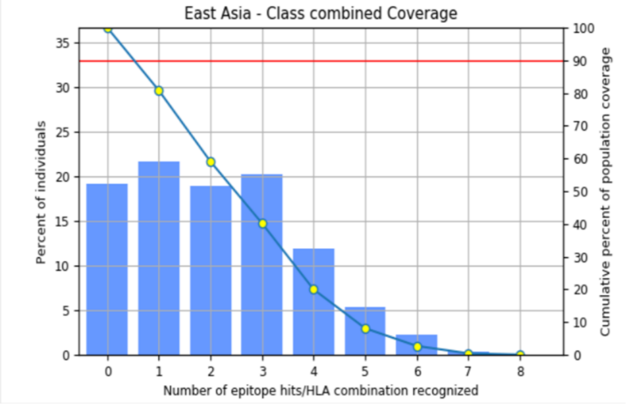


*Population: Europe*

| MHC class | Coverage | Average hit | PC90 |
|---|---|---|---|
| combined | 84.74% | 2.46 | 0.66 |

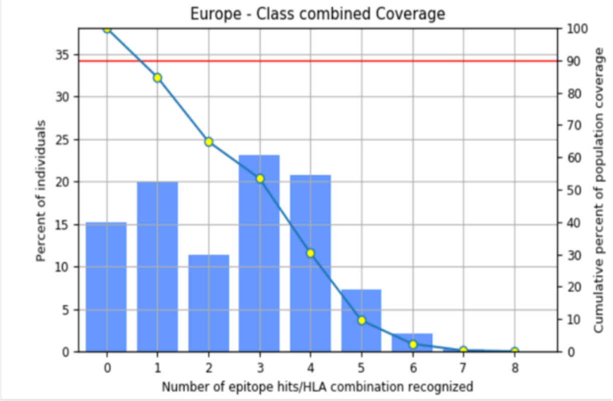


*Population: North Africa*

| MHC class | Coverage | Average hit | PC90 |
|---|---|---|---|
| combined | 77.2% | 2.11 | 0.44 |

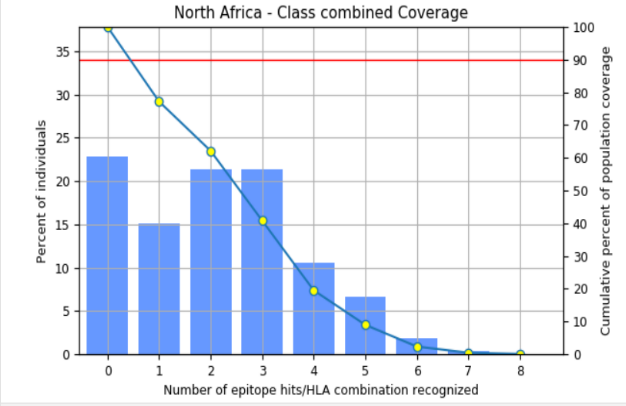

***Population: North America***

| MHC class | Coverage | Average hit | PC90 |
|---|---|---|---|
| combined | 83.87% | 2.53 | 0.62 |

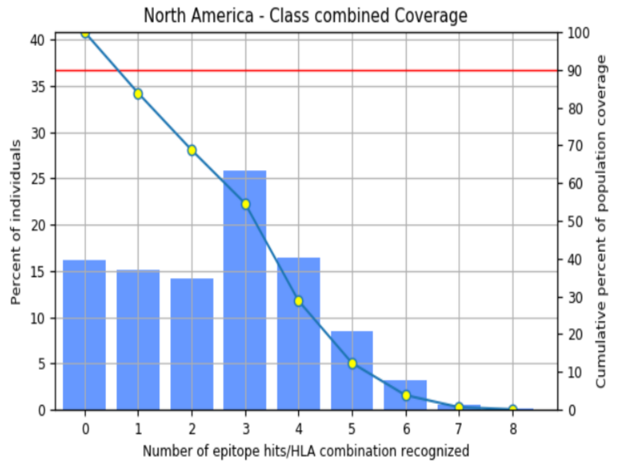


***Population: Northeast Asia***

| MHC class | Coverage | Average hit | PC90 |
|---|---|---|---|
| combined | 63.82% | 1.53 | 0.28 |

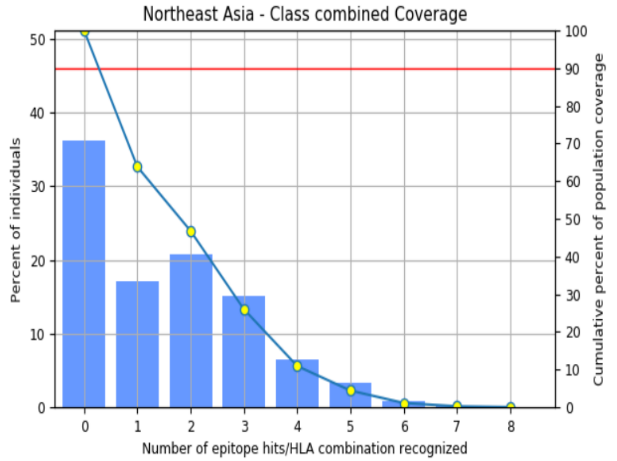


***Population: Oceania***

| MHC class | Coverage | Average hit | PC90 |
|---|---|---|---|
| combined | 34.28% | 0.76 | 0.15 |

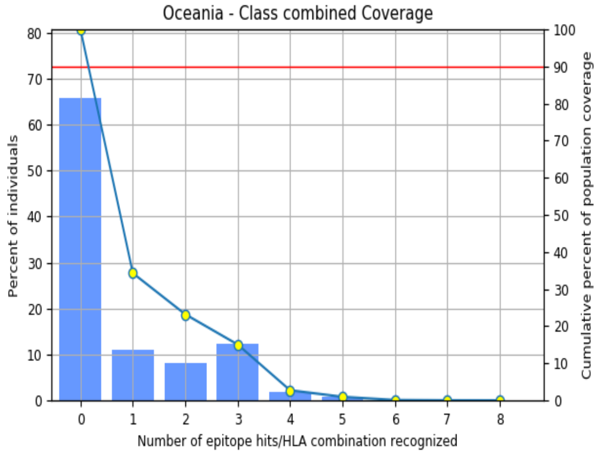


***Population: South Africa***

| MHC class | Coverage | Average hit | PC90 |
|---|---|---|---|
| combined | 66.58% | 1.49 | 0.3 |

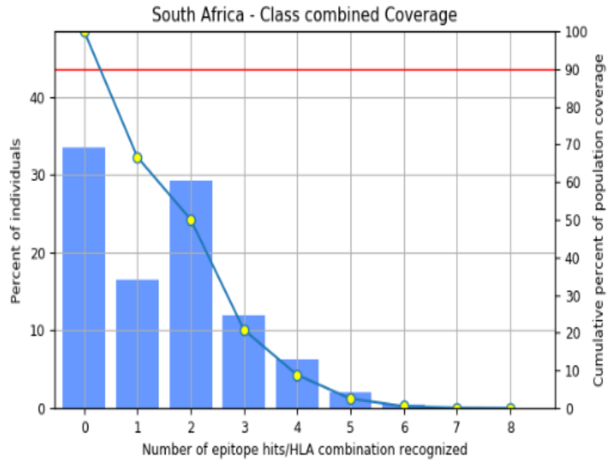


***Population: South America***

| MHC class | Coverage | Average hit | PC90 |
|---|---|---|---|
| combined | 65.0% | 1.6 | 0.29 |

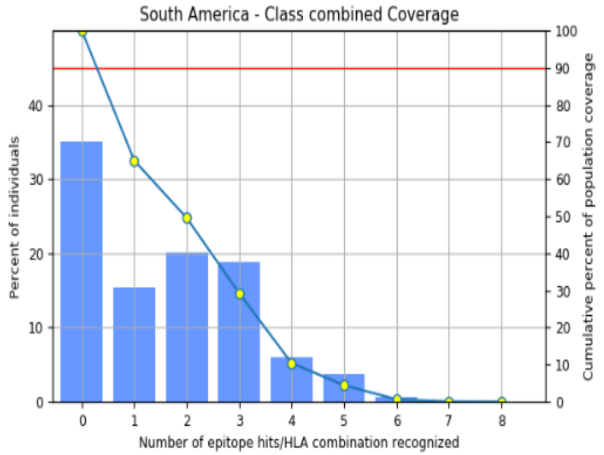


***Population: Southwest Asia***

| MHC class | Coverage | Average hit | PC90 |
|---|---|---|---|
| combined | 71.48% | 1.8 | 0.35 |

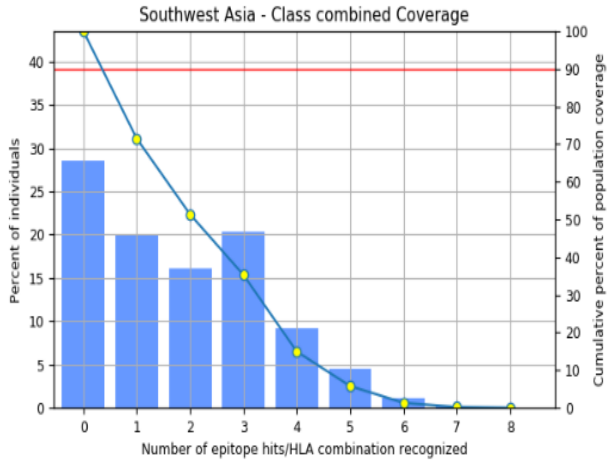

***Population: West Africa***

| MHC class | Coverage | Average hit | PC90 |
| --- | --- | --- | --- |
| combined | 77.0% | 1.99 | 0.43 |

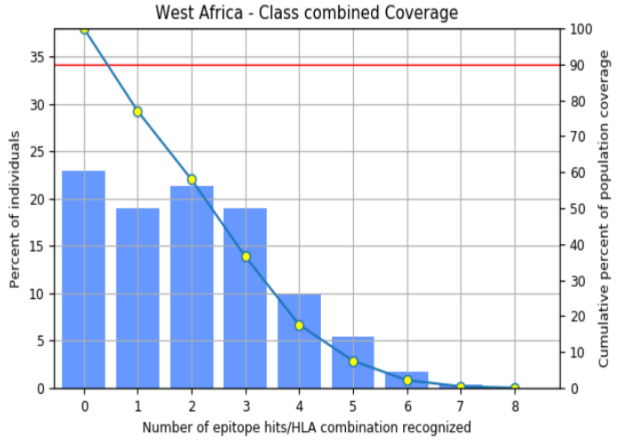


***Population: West Africa***

| MHC class | Coverage | Average hit | PC90 |
| --- | --- | --- | --- |
| combined | 77.0% | 1.99 | 0.43 |

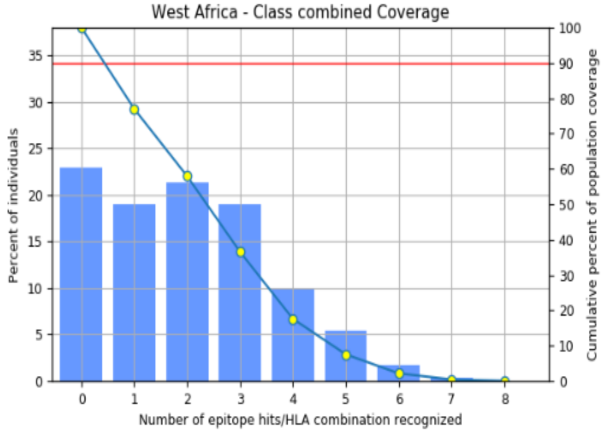


***Population: Southeast Asia***

| MHC class | Coverage | Average hit | PC90 |
| --- | --- | --- | --- |
| combined | 55.76% | 1.37 | 0.23 |

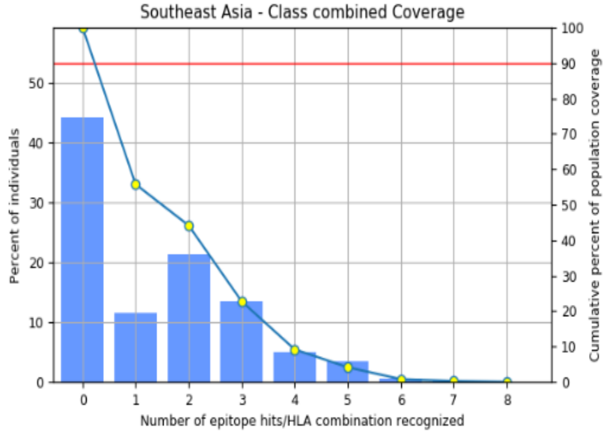


***Population: West Indies***

| MHC class | Coverage | Average hit | PC90 |
| --- | --- | --- | --- |
| combined | 81.52% | 2.29 | 0.54 |

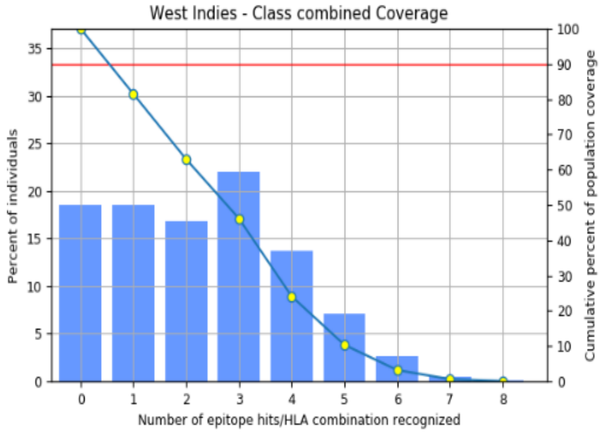


***Population: South Asia***

| MHC class | Coverage | Average hit | PC90 |
| --- | --- | --- | --- |
| combined | 71.46% | 1.64 | 0.35 |

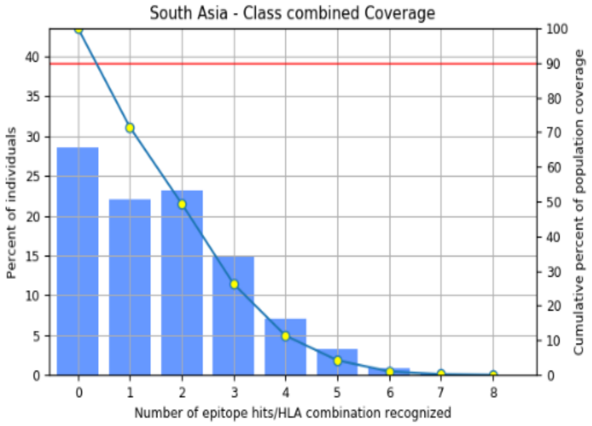
